# A confinable female-lethal population suppression system in the malaria vector, *Anopheles gambiae*

**DOI:** 10.1101/2022.08.30.505861

**Authors:** Andrea L. Smidler, James J. Pai, Reema A. Apte, Héctor M. Sánchez C., Rodrigo M. Corder, Eileen Jeffrey Gutiérrez, Neha Thakre, Igor Antoshechkin, John M. Marshall, Omar S. Akbari

## Abstract

Malaria is among the world’s deadliest diseases, predominantly affecting sub-Saharan Africa, and killing over half a million people annually. Controlling the principal vector, the mosquito *Anopheles gambiae*, as well as other anophelines, is among the most effective methods to control disease spread. Here we develop an innovative genetic population suppression system termed Ifegenia (<u>**I**</u>nherited <u>**F**</u>emale <u>**E**</u>limination by <u>**G**</u>enetically <u>**E**</u>ncoded <u>**N**</u>ucleases to <u>**I**</u>nterrupt Alleles) in this deadly vector. In this bicomponent CRISPR-based approach, we disrupt a female-essential gene, *femaleless* (*fle*), demonstrating complete genetic sexing via heritable daughter gynecide. Moreover, we show that Ifegenia males remain reproductively viable, and can load both *fle* mutations and CRISPR machinery to induce *fle* mutations in subsequent generations, resulting in sustained population suppression. Through modeling, we demonstrate that iterative releases of non-biting Ifegenia males can act as an effective, confinable, controllable, and safe population suppression and elimination system.

## Introduction

Anopheline mosquitoes are responsible for malaria transmission, with the *Anopheles gambiae* complex being the most dangerous, contributing to the quarter billion annual cases (*1*). Controlling anophelines is one of the most effective strategies to prevent disease transmission. However, existing suppression tools such as insecticides and bed nets are becoming increasingly ineffective (*2*). Furthermore, plastic mosquito behaviors may be biasing towards exophagy (*3*), both contributing to the plateau in disease reduction and increasing the costs of control (*1*). Therefore, the development of sustainable, efficient, safe, scalable, and cost-effective vector control technologies is urgently needed.

Because only female mosquitoes transmit disease, most vector control campaigns require exclusive releases of non-biting males. Producing sufficient males *en masse* necessitates development of mechanical, chemical or genetic sex-sorting mechanisms (*4*). Unlike other mosquitoes, sex separation by pupal size is not considered possible in *A. gambiae (5)*. Moreover, lines which permit sex sorting via insecticide resistance are counter reccomended (*6*), and feeding larvae female-killing RNAi yields incomplete phenotypes (*7*), suggesting that transgenic methods may be more robust. Directly sorting males is possible by genetically-encoding fluorescence on sex-chromosomes, near the sex-determinitive loci (*8*), or by using sex-specific alternative splicing or promoters (*9*–*11*). However, these methods require each released male be directly sorted which is labor-intensive, requiring sorting facilities near release sites making releases in remote areas exceedingly difficult. In mosquitoes, genetic approaches that can induce female lethality through removal of chemical inhibition (*12*), or by performing genetic crosses, have been developed (*13*). Unfortunately, these technologies have not been adapted to Anophelines.

In Anophelines, male-biased transgenics have been developed by expressing nucleasess targeting the X chromosome during spermatogenesis (*14*). However these transgenes are constitutively dominant, complicating mass rearing, and are incompletely penetrant, requiring labor-intensive manual selection prior to release. Furthermore, field trials using a precursor system exhibit reduced fitness, presumably due to leaky transgene expression in other tissues targeting the male X (*15*), suggesting an optimal Genetic Sexing System (GSS) should not target male-essential factors. In another sex biasing technology, constitutive overexpression of the male-determining factor, *Yob*, causes male bias in *A. gambiae*, but it is not fully penetrant and faces challenges to scale (*16*). Therefore, while these tools are promising, we agree that the “currently available technology is not scalable” as a GSS for *A. gambiae (4)*. Finally, while sex-distorting gene drives have been developed in *A. gambiae (17)*, these technologies face significant technical, political, ethical, and regulatory hurdles due to the autonomous nature of their spread, making implementation of this technology challenging (*18*). Furthermore, evidence suggests that these types of suppression drives may be hindered by the rapid evolution of resistance alleles (*19*).

To engineer a novel population suppression approach, here we develop a binary CRISPR-based vector control technology targeting a female-essential gene *femaleless* (*fle*) (*20*). Due to its profound daughter-killing phenotypes, we’ve termed it Ifegenia (<u>**I**</u>nherited <u>**F**</u>emale <u>**E**</u>limination by <u>**G**</u>enetically <u>**E**</u>ncoded <u>**N**</u>ucleases to <u>**I**</u>nterrupt Alleles), in honor of Iphigenia of Greek mythology who was sacrificed by her father, King Agamemnon, to win a great battle. Ifegenia operates as a bicomponent approach employing distinct Cas9 and gRNA encoding lines, which result in female-lethality upon hybridization. Offspring incur mosaic somatic, and heritable germline *fle* mutations, resulting in early larval daughter gynecide while leaving sibling males reproductively viable. These males can be iteratively released to load populations with mutations in female-essential gene(s) and causative CRISPR transgenes inducing prolonged, non-driving, population suppression, or sterilized for use in SIT (**Figure S1**). Through modeling, we demonstrate that this technology may be suitable for safe, scalable, confinable, and effective suppression and elimination of *A. gambiae* populations. The technology may also be adapted to other vector species to provide an alternative species-specific population suppression technology to control deadly disease vectors.

## Results

### Engineering strains to target *fle*

We hypothesized that embryonic biallelic CRISPR knockout of *fle* could cause dominant female death. Therefore, to maintain line viability, we developed a bipartite system using separate gRNA and Cas9 lines to induce mosaic *fle* mutations in offspring (**Figure S1**). To ensure robust targeting, we designed and cloned a gRNA-expressing transgene encoding two gRNAs targeting the N-terminal region of *fle*; one targeting 25 bp downstream of the start codon, and a second targeting the first RNA Recognition motif (RRM) (**Figure 1A**). We established three distinct transgenic families (termed gFLE_G_, gFLE_I_, gFLE_J_) by piggyBac-mediated transgenesis and confirmed integration by *Act5C-GFP* selection. For Cas9 expression, we used Vasa2-Cas9 (hereon shortened to Cas9) for its robust germline mutagenesis (**Figure 1A**) (*21*).

**Figure 1.**
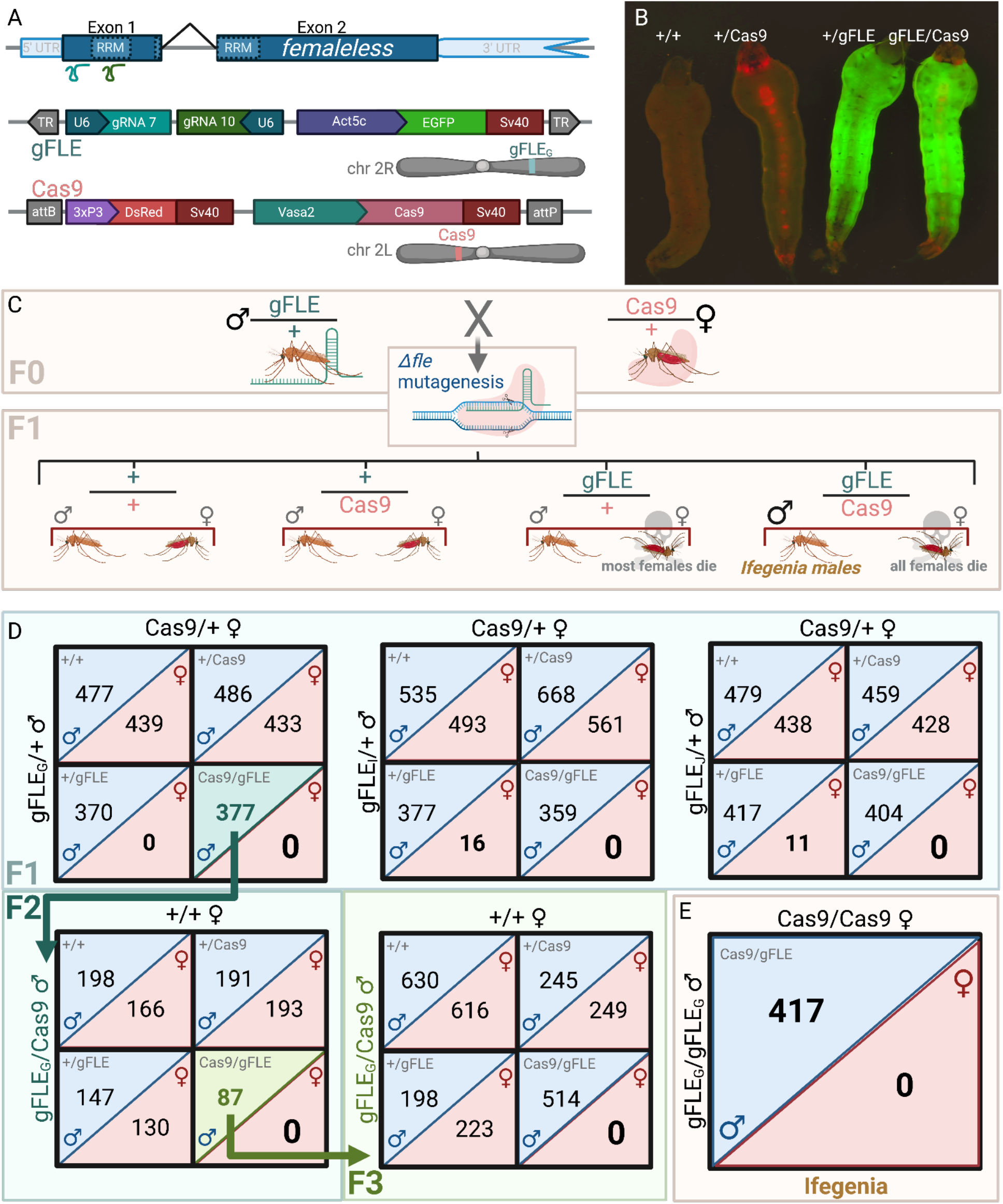
We developed a transgenic crossing system to target *fle* for mosaic knockout. **A)** The gFLE transgene expresses two gRNAs (gRNA7 and gRNA10, teal and emerald respectively) targeting *fle* near the start codon and within the first RNA Recognition Motif (RRM) respectively (gene map, indigo). The PolIII U6 promoter facilitates gRNA expression (Navy), selection is enabled by Act-EGFP-Sv40 (purple, green, red), and the transgene was inserted by piggyBac transgenesis (Terminal Repeats, TR, grey arrows). The Cas9 transgene expresses protein in the adult germline and is maternally deposited into the embryo, and was previously described in Werling et. al, Cell 2019. **B)** Individual genotypes are identified by fluorescence with Cas9 marked by 3xP3-DsRed and gFLE marked by Act-GFP. **C)** Crossing F0 gFLE/+ males to Cas9/+ females yields F1 offspring in Mendelian 1:1:1:1 ratios, with the gFLE/Cas9 female cohort absent. **D)** Absence of all F1 gFLE/Cas9, and some gFLE/+ female offspring, suggests *fle* mutagenesis results in female death. This effect is heritable into the F2 and F3 generations with gFLE_G_/Cas9 males mated to +/+ females able to kill daughters with the appropriate genotypes (F2 and F3 data from gFLE_I_/Cas9 and gFLE_J_/Cas9 males reported in **Figure S7**). **E)** Crossing homozygous gFLE_G_/gFLE_G_ males to Cas9/Cas9 females results in complete elimination of genetic daughters from the progeny. A single gFLE_G_/Cas9 phenotypic daughter was identified, however was genetically male (XY chromosome, data not shown), she was excluded from the analysis (**Table S5**).

### Disrupting *fle* dominantly kills females

To determine the phenotype of gFLE/Cas9 trans-heterozygotes we performed genetic crosses between gFLE-males and Cas9-females and analyzed resulting offspring. Strikingly, all gFLE/Cas9 pupae scored, regardless of family, were phenotypic males (N=638, **Table S1**) indicating robust female death or androgenization before pupation. To confirm *fle* mutagenesis (Δ*fle*), we sequenced gFLE/Cas9 adult males and identified Δ*fle* alleles at both gRNA target sites (**Figure S2**).

To determine if Δ*fle* results in female death, or sufficiently penetrant androgenization indistinguishable from phenotypic males, we crossed gFLE/+ males to Cas9/+ females and quantified the offspring genotypes (**Figure 1B,C**). Strikingly, once again no F1 gFLE/Cas9 adult genetic females were observed in any family during the course of all experiments, and none in the F2 and F3 generations of the gFLE_G_/Cas9 family, despite 2,512 gFLE/Cas9 males reported herein (**Table S1, S2, S14, S17, S18, S19**), and thousands others screened but not reported or quantified. gFLE/Cas9 males (termed as Ifegenia males from hereon) were present at roughly equal numbers to control siblings, suggesting that gFLE/Cas9 females were killed rather than androgenized (**Figure 1D, Table S2**). Supporting this conclusion, PCR confirmed the presence of a Y-chromosome in all randomly selected Ifegenia males (**Figure S3**). Interestingly, all three gFLE families showed a strong reduction in gFLE/+ females, with no gFLE_G_ /+ females identified, and a 23- and 38-fold reduction in females among the gFLE_I_/+ (N = 377:16) and gFLE_J_/+ (N = 417:11) genotypes relative to male siblings respectively (**Figure 1D, Table S2**). This suggests that disrupting *fle* alleles can dominantly kill gFLE/Cas9 inheriting females and that maternal deposition of Cas9 is sufficient for female killing in individuals inheriting gFLE alone.

Because gFLE_G_ yielded the strongest F1 hybrid penetrance among gFLE families, we characterized transgene insertion sites and *fle* mutation profiles of gFLE_G_/Cas9 males by nanopore DNA sequencing, confirming a single transgene insertion and expected *Δfle* alleles (**Figure S4**). To determine relative *fle* expression we performed RNAseq on 3 replicates of 18hr old embryos enriched for the gFLE_G_/Cas9 genotype compared to 3 replicates of each control genotype gFLE_G_, Cas9 and WT. We found a modest but highly significant reduction in *Fle* (padj ranging from 4E-06 -2E-08) (**Tables S3-S10)** with non-mutant genotypes clustering together as expected (**Figure S5**).

To better understand the effect of *fle* downregulation, we performed gene ontology (GO) analysis on gFLE_G_/Cas9 vs control genotypes revealing expected upregulation of DNA-repair and mitosis/cell-cycle arrest-related terms, and somewhat unexpected downregulation of mitochondria and protein metabolism (**Tables S11 and S12**). Taken together, these results indicate that we are indeed disrupting the gene *fle* which is resulting in observable Δ*fle* mutations in the DNA and reduced *fle* transcript expression leading to transcriptome wide effects.

### *Fle* is essential for female larval development

Next, to determine the life stage of female death we quantified the genotypes of one-day old larvae from the cross outlined in **Figure 1C**, and did not observe a significant reduction in gFLE/Cas9 or gFLE/+ genotypes, indicating that the majority of gFLE/Cas9 females survive embryogenesis (**Figure S6, Table S13**). We then determined that most gFLE/Cas9 female death occured during the larval stage by monitoring and quantifying the survival of each sex-genotype from hatching through pupation (**Figure S7, Table S14**). Together, our results demonstrate that CRISPR-targeting of *fle* has remarkable larval female-killing penetrance.

### Ifegenia males are reproductively viable

To be candidates for release in population suppression campaigns Ifegenia adult males must be long lived, and reproductively viable. To quantify adult longevity, we monitored gFLE_G_/Cas9 and gFLE_J_/Cas9 males compared to +/+ siblings in survival assays. Male gFLE_G_/Cas9, but not gFLE_J_/Cas9, displayed slight reductions in longevity (**Figure S8, Table S15**). Notwithstanding, because males mate between 4 and 8 days old, the effects of this lifespan reduction should be minimal and surmountable by additional releases (*22, 23*). To verify reproductive viability, we crossed Ifegenia males to wild type females and confirmed the presence of viable progeny (**Table S16**). Though more formal characterization of male fitness was beyond the scope of this work, taken together these results suggest that Ifegenia males should be sufficiently reproductively viable to achieve population suppression following iterative releases, however large-scale cage and field trials should be undertaken in the future.

### Ifegenia induces genetic sexing

Given the profound female-killing properties of Ifegenia, we sought to determine its penetrance of genetic sexing for use in male-only vector control campaigns (**Figure S1**). We mated homozygous gFLE_G_/gFLE_G_ males to Cas9/Cas9 females and verified the complete absence of females among all offspring (**Figure 1E, Table S17**). As expected, complete killing of genetic females was observed (N = 417, gFLE_G_/Cas9 males). Interestingly however, we identified a single, likely sterile, phenotypic gFLE_G_/Cas9 female as defined by the pupal genitalia parameters of our assay. This individual PCR-amplified for the Y-chromosome revealing a rare male-feminization intersex genotype (**Table S17**). Taken together, this reveals a powerful application of Ifegenia as a tool for genetic sexing of *A. gambiae* (**Figure S1**).

### Ifegenia males confer multi-generational daughter gynecide

We next sought to determine if Ifegenia males can ‘load’ non-driving CRISPR transgenes and Δ*fle* alleles into subsequent generations (F2 and F3), and cause multi-generational daughter-killing effects (**Figures 1D, S2, S9**). We observed approximately normal Mendelian inheritance of transgenes in all families, and complete female-killing of all F2 and F3 gFLE_G_/Cas9 females (**Figure 1D, Tables S18, S19**). Moreover, out of the few F2 and F3 gFLE/Cas9 females observed from families gFLE_I_ and gFLE_J_, all either died as pupae or young adults, appeared androgenized, or were feminized XY genetic males. We also determined the approximate frequency of Δ*fle* alleles in the F2 generation by PCR and enzyme digest (*24*). We observed 85-95% *Δfle* allele frequency among F2 individuals inheriting one or no transgenes, representing a conservative estimate on the frequency of germ cell mutagenesis in the parental F1 gFLE_G_/Cas9 germline. Importantly, this work indicates that *fle* is haplosufficient, in contrast to earlier hypotheses (*20*). Among gFLE_G_/Cas9 F2 progeny, 100% harbored at least one *Δfle* allele, however this is likely inflated due to active lethal mosaicism occurring in this genotype (**Figure S10**)(*13, 25*). Together these results indicate that Ifegenia males are reproductively viable, and pass on both CRISPR transgenes and *Δfle* alleles to subsequent generations resulting in multi-generational daughter gynecide.

### IIfegenia induces confinable population suppression

To determine whether iterative releases of fertile Ifegenia males could facilitate population suppression and elimination, we incorporated the above data on Ifegenia performance into a mathematical model (*26*) and simulated releases into a population of 10,000 *A. gambiae* adults (**Figure 2**). Weekly releases of up to 500 Ifegenia eggs (female and male) per wild-type adult (female and male) were simulated over 1-48 weeks. The scale of these releases was chosen considering adult release ratios of 10:1 are common for sterile male mosquito interventions (*27*) and female *A. gambiae* produce >30 eggs per day in temperate climates (*28*). We considered Ifegenia constructs with 1-3 gRNA target sites, and given the impressive laboratory results, simulated target site cutting rates of 90-100% per allele and maternal deposition of Cas9 whenever expressed by the mother. Male mating competitiveness and female fecundity were assumed to not be impacted by the constructs. Results from these simulations suggest that significant population suppression (>90%) that endures for >2 years is observed for a wide range of achievable release schemes, e.g., 22 weekly releases of 300 or more Ifegenia eggs per wild adult, and elimination is expected to occur in >90% of simulations for 26 weekly releases of the same size. Interestingly, Ifegenia performance is robust to several system features -population suppression does not significantly differ as target site cutting rates vary from 90-100%, or as the number of gRNA target sites increase from 1-3. These findings suggest that Ifegenia can achieve robust temporary population suppression over a wide range of release parameters, however permitting rebound of native populations after ceasing releases. Opportunity for population rebound may be desired in locales where disease elimination has been achieved, and where ecological concerns warrant return of the native mosquito, making Ifegenia a valuable vector control technique for the toolkit.

**Figure 2.**
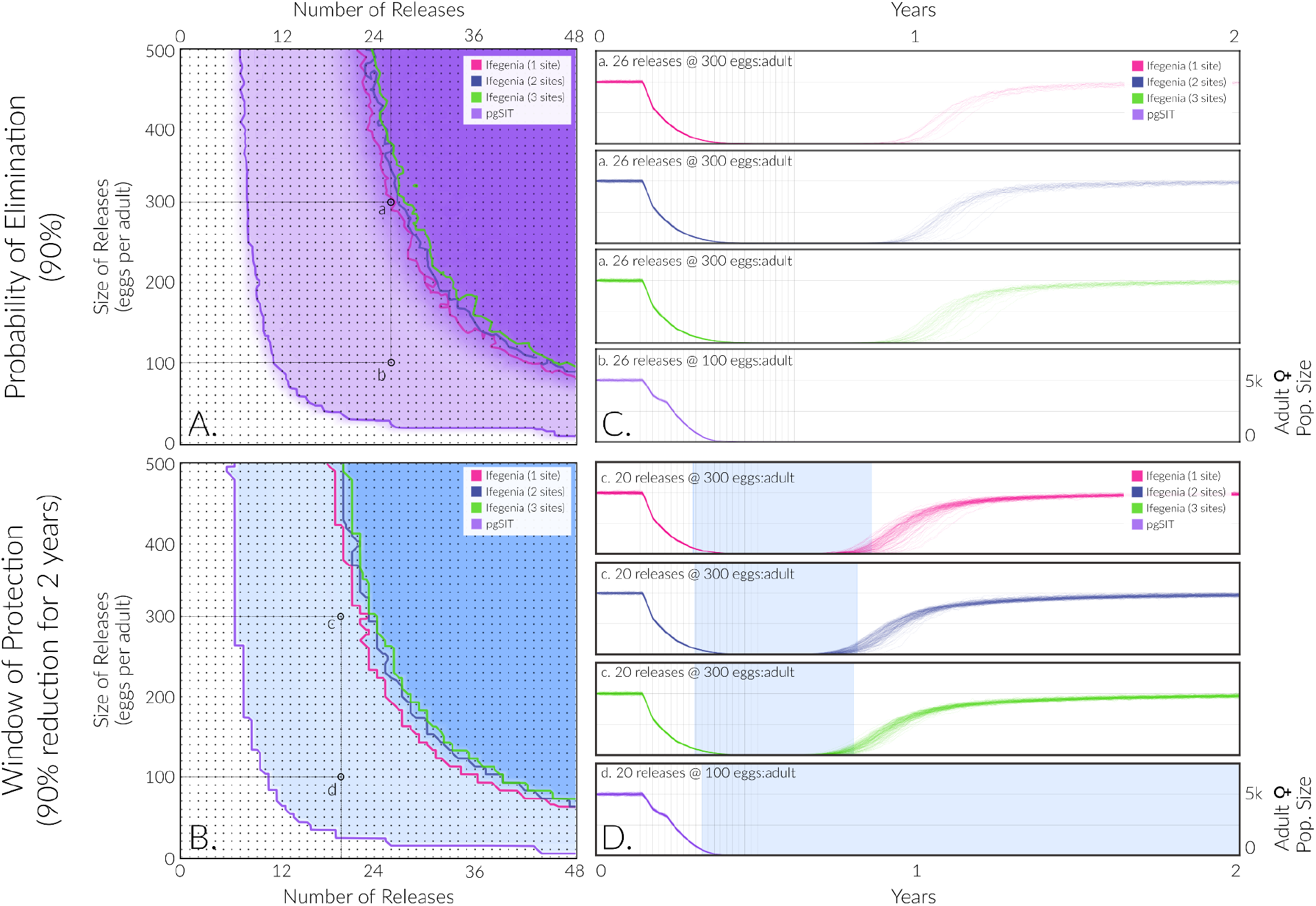
Model-predicted impact of releases of Ifegenia (1-3 target sites) and pgSIT eggs on *A. gambiae* population density and elimination. Weekly releases were simulated in a randomly-mixing population of 10,000 adult mosquitoes using the MGDrivE simulation framework (*1*) and parameters described in **Table S20**. Weekly releases of up to 500 Ifegenia or pgSIT eggs were simulated over 1-48 weeks. Ifegenia (1-3 target sites) and pgSIT were simulated with cutting rates of 100% per allele and maternal deposition of the Cas whenever expressed by the mother. Male mating competitiveness and female fecundity were not impacted by the constructs. For Ifegenia, females homozygous for any Δ*fle* mutant allele were considered unviable, and for pgSIT, males having the system were considered sterile. Elimination probability **(A)** was calculated as the percentage of 120 stochastic simulations that resulted in *A. gambiae* elimination, for each parameter set. Window of protection **(B)** was calculated as the percentage of 120 stochastic simulations that resulted in the *A. gambiae* population being suppressed by at least 90% for at least two years, for each parameter set. In both cases, contours represent thresholds for each metric and system -i.e., parameter sets for which 90% of simulations result in elimination, or for which simulations result in a median window of protection of at least two years. Sample time-series depicting adult female *A. gambiae* population density are depicted on the right demonstrating elimination **(C)** and window of protection **(D)**. For window of protection, the shaded region represents the time during which the median adult female population density is reduced by 90%. Population elimination and an extended window or protection are possible for a wide range of release scenarios for each system. For Ifegenia, elimination is achieved ∼90% of the time for >26 releases at a release ratio of ∼300:1 eggs per wild adult, and a median window of protection >2 years is achieved for >22 releases at the same ratio. Interestingly, the number of gRNA target sites does not significantly impact these results. Ifegenia can achieve similar results to pgSIT, although it requires slightly more and/or larger releases.

## Discussion

Historically establishing scalable genetic sexing systems in anophelines has proven problematic, with many lines incompletely penetrant (*14, 16*), and most relying on fluorescence-based, or manual sorting, directly prior to release (*11, 15*). Inducible genetic systems which automatically kill the undesired sex have been largely unattainable due to limited understanding of the sex-determination pathway (*29*), and husbandry limitations which prohibit development of translocation lines (*4*).

In this work we demonstrate that targeting *fle* in *A. gambiae* results in complete killing of genetic females through Lethal Mosaicism (*13, 25*) without severely curbing male fitness, and develop Ifegenia as a novel non-driving intervention for prolonged population suppression. Ifegenia’s bicomponent design provides three main advantages over alternative technologies. i) It is ideal for industrial mass production, as the two homozygous stock strains exhibit normal sex ratios and fertility with only F1s displaying female-killing, providing the most powerful genetic sexing mechanism in anophelines to date. ii) This design enables direct release of F1 eggs/larvae/pupae into the wild, eliminating wasted resources on rearing females, and obviating injurious manual sorting (*4*), making possible a myriad of scalable, cost-effective, distribution modalities. And iii) because Cas9 and gRNAs are isolated until the hybridization cross, creation of CRISPR-generated resistance alleles in stock lines is impossible, preventing roadblocks faced by many gene drives (*17, 30*).

Currently scalability is only limited by the manual sex separation required to establish F0 parental crosses between gFLE and Cas9 stock lines. However, because a single gFLE-fertilized Cas9 female can conservatively yield 200 pre-sexed Ifegenia sons over her lifetime this provides an order-of-magnitude efficiency improvement compared to alternatives (**Figure S1**). Future iterations of Ifegenia could be adapted to include sex-specific fluorophores to automate F0 parental sex sorting by COPAS, further improving scalability, and streamlining mass rearing and releases on a continental scale (*8–11, 31*)

Our modeling suggests iterative releases of fertile Ifegenia males can load populations with female-killing alleles, and causative CRISPR transgenes, ensuring prolonged population suppression effects without drive. This however would require iterative releases and temporary persistence of GM mosquitoes, and could concurrently generate resistance alleles in wild populations, potentially hindering future campaigns targeting these loci. Importantly however, generation of fully resistant alleles is attenuated by targeting two distinct loci within *fle*, and in the future can be improved by multiplexed targeting of additional female-essential genes, which modeling suggests should achieve similar performance for population suppression. Taken together, we believe Ifegenia fills a valuable population suppression niche as an intermediate between technologies such as pgSIT (*13*) and suppression gene drives (*17, 32*).

Importantly, this work demonstrates *fle*’s potential to enable novel GM vector control tools beyond those outlined here. Due to conservation among anophelines, development of Ifegenia in related malarial species such as *A. arabiensis, A. colutzii*, and *A. stephensi* should be trivial. Additionally, manipulation of *Fle*, could enable a novel suite of tet-inducible/tet-supressible sex separation systems, homing based gene drives (*33*–*35*), and Y-drives (*36*), among other technologies.

The magnitude of effects insecticides have on non-target populations is hard to understate, given their probable culpability behind the decline in global insect biomass, and the disastrous cascading multi-trophic ecosystem perturbations which result (*37*). In 2019 alone, over 25 billion m^2^ of insecticides were applied globally, the majority of which targeting malaria vectors, making the ecological impacts on Africa and Asia particularly harmful (*38*). Due to the decreasing efficacy of insecticides and the risk of further damaging ecosystems, alternative vector control strategies, like Ifegenia, must be explored and implemented to revolutionize control of these deadly species.

## Materials and Methods

### Mosquito rearing and maintenance

*A. gambiae* was derived from the G3 strain. The mosquitoes were reared in 12hr light/ dark cycles at 27°C with 50-80% humidity in cages (Bugdorm, 24.5 × 24.5 × 24.5 cm) in an ACL-2 insectary. Adults were provided with 0.3 M aqueous sucrose *ad libitum*, and females were blood fed on anesthetized mice for 2 consecutive days for ∼15 min at a time. Males and females were allowed to mate for at least 2 days prior to a blood meal. Egg dishes were provided 2 days after a blood meal. Eggs were allowed to melanize for 2 days before being floated in trays. Larvae were reared and fed, as well as pupae screened and sexed, in accordance with established protocols,, (*39*).

### gRNA Design, cloning and transgenesis

The *Fle* (AGAP013051) target gene reference sequence was extracted from VectorBase (*40*). To verify target sequence and detect any polymorphisms, gDNA was extracted (Qiagen, DNeasy Blood & Tissue Kits, Cat. No. / ID: 69504) from pools of 10 individuals, cloned into pJET (ThermoFisher Scientific, Cat. No./ID: K1231), and individual colonies were sanger sequenced for the *fle* locus. Putative candidate gRNAs in conserved regions were identified using http://crispor.tefor.net/ two gRNAs, gRNA7 [5’-CGACGGCTCGTTCATCGCTG**GGG**-3’] and gRNA10 [5’-ATCGAGCGCGTCGCCTGGTA**CGG-**3’] were selected. Each gRNA targets Exon 1, and overlaps a semi-unique restriction enzyme site to facilitate downstream screening and identification of mutant alleles (*24*). We modified a piggyBac transgenesis backbone pbVTKactR (*41*) to contain each gRNA under expression of the *A. gambiae* U6 promoter (*21*) synthesized as gBlocks®, and replaced endogenous Red for m2Turquoise (ECFP) The final plasmid sequence (plasmid 1154B, transgene gFLE) was confirmed by Sanger sequencing and is available on Addgene (#187238). Embryonic microinjections of gFLE into G3 wildtype embryos was carried out as described previously (*31*). We identified 9 transgenic founders, which were individually outcrossed to wild type to establish distinct families. Families gFLE_G_, gFLE_I_, and gFLE_J_ were selected for analysis.

### Fluorescent Sorting, Sexing and Imaging

*A. gambiae* were fluorescently sorted, sexed, and imaged using the Leica M165FC fluorescent stereomicroscope using a Leica DMC2900 camera. Fluorescence was visualized using the CFP/YFP/mCherry triple filter, and was sexed by examination of pupal genital terminalia. In cases where sex was indeterminable by pupal phenotype, genotype was validated by Y-chromosome PCR (see below).

### Genetic Cross Setup

For all crosses, pupae were fluorescently sorted and sexed, and allowed to emerge as adults in separate cages to ensure female virginity before crossing. Unless otherwise indicated, crosses were set up with 1-3 day old adults, allowed to mate *ad libitum* for 4 days, then blood fed. For preliminary test crosses, 50 gFLE-positive males and 50-Cas9 positive females (mixed heterozygotes and homozygotes) were crossed, and offspring pupal sex-genotypes were scored (**Table S1**). For crosses requiring heterozygous parents, gFLE-positive or Cas9-positive individuals were first crossed to wild type of the reciprocal sex and fluorescently sorted to generate guaranteed heterozygous F0’s. These F0 +/gFLE males and +/Cas9 females were then intercrossed for analysis of the Mendelian inheritance patterns in the F1 offspring. For assays requiring homozygous X homozygous mating pairs, one of two assays was set up. In early experiments, small cages of gFLE-positive males (enriched for homozygotes by fluorescence intensity) were mated to pure Cas9/Cas9 females, allowed to mate *en masse* and allowed to oviposit *en masse*. Only those broods which yielded 100% gFLE/Cas9 offspring were considered from gFLE/gFLE X Cas9/Cas9, and scored. In a second experiment, larger cross cages were set up essentially as described above, however females were isolated into iso-female ovicups prior to oviposition. Individual broods were screened for pure hybrid transheterozygous (gFLE/Cas9) offspring, and only those with this genotype were scored.

### Embryo survival assays

To determine if females were dying during embryogenesis, +/gFLE males and +/Cas9 were crossed. From large egg lays, a random sampling of ∼500 unhatched embryos were separated from the egg dish and allowed to hatch. All larvae were scored by genotype at 1 day post-hatching and reported in **Figure S6, Table S13**.

### Larvae survival

To determine if females were dying as larvae (after hatching and before pupation), we fluorescently sorted 1do larval offspring of the +/gFLE x +/Cas9 cross. 40 larvae from each genotype were reared in separate trays, then scored as pupae by sex and genotype and are reported in **Figure S5 and Table S14**.

### *Fle* knockout mutant analysis

DNA was individually extracted from L3-L4 *A. gambiae* larvae using the DNeasy Blood & Tissue Kits (Qiagen, Cat. No. / ID: 69504). 1 µl of genomic DNA was used as a template in a 20 µl PCR reaction using Q5 HotStart DNA polymerase (NEB, Cat. No./ID: M0493L) and primers 1154A.S5 and 1154A.S7 amplifying genomic *Fle* sequences. The resulting product was run on a 1% agarose gel in TAE buffer), gel extracted with the Zymoclean Gel DNA Recovery Kit (Zymo Research, Cat. No./ID: D4007), cloned into the pJET vector (Thermo Scientific, Cat. No. / ID: K1231), transformed into chemically competent *E. coli* (Promega, JM109), and plated on LB-Ampicillin plates. Sanger sequencing reads from individual colonies represented amplicons from individual mutant alleles, (primers PJET1-2F and/or PJET 1-2R) were compared to *fle* sequences from our WT mosquitoes as a reference genome, and a selection are summarized in **Figure S2**. All primer sequences can be found in **Table S21**.

### *Δfle* allele quantification

To quantify *Δfle* mutant alleles, we performed PCR amplification followed by direct digest on *Fle* amplicons in F2 individuals. Genomic DNA samples were prepared using the Qiagen DNeasy extraction kit (Cat. No. / ID: 69504), and amplified in 50 µl PCR reactions using Taq DNA polymerase (NEB, Cat. No. / ID: #M0273S) and primers 1154A.S8 and 1154A.S29 (1121bp). PCR product was divided into three 15 µl aliquots; one was undigested for reference, one digested with BstNI (Cat. No. / ID: R0168S), and one with BseYI (NEB cat# R0635S) according to manufacturer’s protocols. PCR amplicons from WT alleles are expected to digest into 228 bp and 893 bp, and 403 bp and 718 bp fragments for BseYI (gRNA7) and BstNI (gRNA10) respectively.

Failure to digest a significant quantity of PCR product indicates a likely polymorphism under the gRNA target site. Immediately after digestion, the 15ul raw PCR product, 15ul BstNI-digested product, and 15ul BseYI-digested product were run side-by-side on gel (1% agarose gel in TAE buffer, 1 kb ladder (NEB, Cat. No. / ID: N3232L), most gels run at 115V for 40 minutes). Gel images shown in **Figure S10**.

### Y chromosome amplification

Samples were extracted with the Qiagen DNeasy extraction kit (Cat. No. / ID: 69504). 1ul of genomic DNA was used as a template in a 20ul PCR reaction using Q5 HotStart DNA polymerase (NEB, Cat. No./ID: M0493L). Each individual was genotyped for the presence of the y-chromosome (*42*). Positive control PCRs used *A. gambiae*-specific 28S primers (1123A.S2/1123A.S3, 230 bp fragment) (*43*).

### Mathematical modeling

To model the expected performance of Ifegenia and pgSIT at suppressing and eliminating local *A. gambiae* populations, we used the MGDrivE simulation framework (*26*). This framework models the egg, larval, pupal, and adult mosquito life stages with overlapping generations, larval mortality increasing with larval density, and a mating structure in which females retain the genetic material of the adult male with whom they mate (sperm) for the duration of their adult lifespan. The inheritance patterns of the Ifegenia and pgSIT systems were modeled within the inheritance module of MGDrivE. For simplicity, the inheritance model assumes that mosquitoes having the Cas9 allele and a gRNA targeting the N-terminal region of *fle*, or targeting male fertility in the case of pgSIT, cleave recessive terminal regions of *fle* and/or of the male fertility allele according to a defined frequency. In the case of multiple target sites, cleavage of each target allele is treated as an independent event. For Ifegenia, females homozygous for any *fle* mutant allele have reduced viability, and in the case of pgSIT, male mosquitoes homozygous for the fertility mutant allele have reduced fertility. A proportion of progeny of female mosquitoes having the gRNA and Cas9 alleles also have their N-terminal region of *fle* or, in the case of pgSIT, male fertility allele, cleaved due to the maternal deposition of Cas9. For this analysis, it is assumed that there are no male mating competitiveness or other fitness costs associated with having the gRNA or Cas9 alleles, and that individuals heterozygous for either allele are not affected. The inheritance patterns in this model did not account for the development of resistance alleles, which could potentially inhibit the cleavage of gRNA target sites.

We considered a single randomly-mixing *A. gambiae* population of 10,000 mosquitoes and implemented the stochastic version of the MGDrivE framework to capture random effects at low population sizes and the potential for population elimination. Density-independent mortality rates for juvenile life stages were calculated for consistency with the population growth rate in the absence of density-dependent mortality, and density-dependent mortality was applied to the larval stage following Deredec *et al*. (*44*). Weekly releases of up to 500 transgenic eggs per wild adult mosquito (female and male) were simulated over a period of 1-48 weeks. The scale of egg releases was chosen following the precedent in Kandul *et al. (25)* for equivalence to an adult release ratio on the order of 10:1, taking into account survival of released eggs through to the adult life stage in the presence of density-dependent larval mortality. 120 repetitions were carried out for each parameter set, and mosquito genotype trajectories, along with the proportion of simulations that led to local population elimination, were recorded. Complete model and intervention parameters are listed in **Table S20**.

### Determination of transgene integration sites

To determine the transgene insertion sites, we performed Oxford Nanopore genome DNA sequencing. We extracted genomic DNA using the Blood & Cell Culture DNA Midi Kit (Qiagen, Cat. No. / ID: 13343) from 16 adult Ifegenia transheterozygous males harboring both transgenes (Cas9/+ ; gFLE /+), following the manufacturer’s protocol. The sequencing library was prepared using the Oxford Nanopore SQK-LSK110 genomic library kit and sequenced on a single MinION flowcell (R9.4.1) for 72 hrs. Basecalling was performed with ONT Guppy base calling software version 6.1.2 using dna_r9.4.1_450bps_sup model generating 3.02 million reads above the quality threshold of Q≧ 10 with N50 of 12308 bp and total yield of 19.49 Gb. To identify transgene insertion sites, nanopore reads were aligned to plasmids carrying either gFLE (1154B, Addgene as plasmid #187238) or Cas9 (*21*) constructs using minimap2 (*45*). Reads mapped to the plasmids were extracted and mapped to the *A. gambiae* genome (GCF_000005575.2_AgamP3). Exact insertion sites were determined by examining read alignments in Interactive Genomics Viewer (IGV). The gFLE_G_ transgene is integrated between positions 23,279,556 and 23,279,559 on chromosome 2R (NT_078266.2) and falls into the last intron of AGAP002582. Cas9 is inserted between positions 10,326,500 and 10,326,503 on 2L (NT_078265.2). The site is located in the intergenic region between AGAP005126 and AGAP005127 as expected by its integration in the X1 docking site (*31*). Using nanopore data, we also confirmed genomic deletions in the target gene, AGAP013051, as expected (**Figure S4)**. The nanopore sequencing data has been deposited to the NCBI sequence read archive (PRJNA862928, reviewer link: https://dataview.ncbi.nlm.nih.gov/object/PRJNA862928?reviewer=pbmvm5fb9in78ro2a1e7p227 oc)

### Transcriptional profiling and expression analysis

To quantify target gene reduction and expression from transgenes as well as to assess global expression patterns, we performed Illumina RNA sequencing. A cross of 50 heavily enriched gFLE_G_/gFLE_G_ males to 50 homozygous Cas9/Cas9 females was performed, allowed to mate *ad libitum* for 5 days, and blood fed. 72 hours after the blood feeding, the egg dish is placed into cages for synchronous egg laying, and females allowed to oviposit for 2 hours. Eggs were collected 18 hours after the first egg lay was observed. We extracted total RNA using miRNeasy Tissue/Cells Advanced Mini Kit (Qiagen, Cat. No. / ID: 217604) from 100uL embryos, estimate volume, with each genotype (WT; +/Cas9; +/gFle; gFle/Cas9) in biological triplicate (12 samples total), following the manufacturer’s protocol. Genomic DNA was depleted using the gDNA eliminator column provided by the kit. RNA integrity was assessed using the RNA 6000 Pico Kit for Bioanalyzer (Agilent Technologies, Cat. No. / ID: #5067-1513), and mRNA was isolated from ∼1 μg of total RNA using NEBNext Poly(A) mRNA Magnetic Isolation Module (NEB, Cat. No. / ID: E7490). RNA-seq libraries were constructed using the NEBNext Ultra II RNA Library Prep Kit for Illumina (NEB, Cat. No. / ID: E7770) following the manufacturer’s protocols. Briefly, mRNA was fragmented to an average size of 200 nt by incubating at 94°C for 15 min in the first strand buffer. cDNA was then synthesized using random primers and ProtoScript II Reverse Transcriptase followed by second strand synthesis using NEB Second Strand Synthesis Enzyme Mix. Resulting DNA fragments were end-repaired, dA tailed, and ligated to NEBNext hairpin adaptors (NEB, Cat. No. / ID: E7335). Following ligation, adaptors were converted to the “Y” shape by treating with USER enzyme, and DNA fragments were size selected using Agencourt AMPure XP beads (Beckman Coulter #A63880) to generate fragment sizes between 250-350 bp. Adaptor-ligated DNA was PCR amplified followed by AMPure XP bead clean up. Libraries were quantified using a Qubit dsDNA HS Kit (ThermoFisher Scientific, Cat. No. / ID: Q32854), and the size distribution was confirmed using a High Sensitivity DNA Kit for Bioanalyzer (Agilent Technologies, Cat. No. / ID: 5067-4626). Libraries were sequenced on an Illumina NextSeq2000 in paired end mode with the read length of 50 nt and sequencing depth of 20 million reads per library. Base calls and FASTQ generation were performed with DRAGEN 3.8.4. The reads were mapped to the VectorBase-58_AgambiaePEST genome supplemented with gFLE and Cas9 transgene sequences using STAR. On average, ∼97.4% of the reads were mapped (**Table S7**). Gene expression was then quantified using featureCounts against VectorBase annotation release 58 GTF (VectorBase-58_AgambiaePEST.gtf). TPM values were calculated from counts produced by featureCounts and combined (**Table S8**). Hierarchical clustering of the data shows that for each genotype, all replicates cluster together, as expected (**Figure S10**). DESeq2 was then used to perform differential expression analyses between controls (WT; +/Cas9; +/gFle) and gFle/Cas9 (**Table S9-S13**). For each DESeq2 comparison, gene ontology enrichments were performed on significantly differentially expressed genes using R package topGO. Illumina RNA sequencing data has been deposited to the NCBI-SRA, (PRJNA862928, reviewer link: https://dataview.ncbi.nlm.nih.gov/object/PRJNA862928?reviewer=pbmvm5fb9in78ro2a1e7p227 oc)

### Statistical analysis

Statistical analysis was performed in JMP8.0.2 by SAS Institute Inc and Prism9 for macOS by GraphPad Software, LLC. At least three biological replicates were used to generate statistical means for comparisons. P values were calculated for a two-sided Student’s t-test with equal or unequal variance. A two-sided F test was used to assess the variance equality. The departure significance for survival curves was assessed with the Log-rank (Mantel-Cox) and Gehan-Breslow-Wilcoxon texts. Multiple comparisons were corrected by the Bonferroni method. All plots were constructed using Prism 9.1 for macOS by GraphPad Software, LLC.

### Ethical conduct of research

All animals were handled in accordance with the Guide for the Care and Use of Laboratory Animals as recommended by the National Institutes of Health and approved by the UCSD Institutional Animal Care and Use Committee (IACUC, Animal Use Protocol #S17187) and UCSD Biological Use Authorization (BUA #R2401).

### Data availability

Complete sequence maps and plasmids are deposited at Addgene.org (#187238). All Illumina and Nanopore sequencing data has been deposited to the NCBI-SRA PRJNA862928, reviewer link: https://dataview.ncbi.nlm.nih.gov/object/PRJNA862928?reviewer=pbmvm5fb9in78ro2a1e7p227 oc. All data used to generate figures are provided in the Supplementary Materials/Tables. Generated *A. gambiae* transgenic lines are available upon request to O.S.A.

## Acknowledgements

We thank Judy Ishikawa and Michelle Bui for help with mosquito husbandry, Dhara Desai, Sanle Chen, Akshay Bharadwaj, and Martha L. Chow for laboratory assistance. Figures were Created with BioRender.com. This work was supported by funding from a DARPA Safe Genes Program Grant (HR0011-17-2-0047) and an NIH award (R01AI151004) awarded to O.S.A., and by funds from the Bill & Melinda Gates Foundation (INV-017683) awarded to J.M.M. The views, opinions, and/or findings expressed are those of the authors and should not be interpreted as representing the official views or policies of the U.S. government.

## Author contributions

O.S.A and A.L.S conceived and designed the experiments. J.P., R.A., A.L.S. and N.T. performed molecular and genetic experiments as well as mosquito crosses and husbandry I.A. performed the Oxford nanopore and RNA sequencing experiments and analysis. H.M.S.C., R.M.C., E.J.G. and J.M.M. performed mathematical modeling. All authors contributed to the writing, analyzed the data, and approved the final manuscript.

## SUPPLEMENTARY TABLES

**Table S1:** Preliminary female-killing crosses. Raw sex-genotype pupae count for **F1** offspring from [+/gFLE & +/gFLE/gFLE] X [Cas9/Cas9 & +/Cas9] crosses.

**Table S2:** Raw sex-genotype pupae count for **F1** offspring from +/gFLE X +/Cas9 crosses. (Presented in **Fig 1D** Top row).

**Table S3**: RNAseq Mapping Stats for all 12 samples.

**Table S4**: Combined RNAseq TPM values for all 12 samples.

**Table S5**: RNAseq differential expression analysis comparing Vasa-*Cas9* vs. WT -control

**Table S6**: RNAseq differential expression analysis comparing gFLE(G) vs. WT

**Table S7**: RNAseq differential expression analysis comparing Vasa-*Cas9* vs. gFLE(G) transheterozygotes -control

**Table S8**: RNAseq differential expression analysis comparing Vasa-*Cas9 /*gFLE(G) transheterozygotes vs. WT

**Table S9**: RNAseq differential expression analysis comparing Vasa-*Cas9 /*gFLE(G) transheterozygotes vs. Vasa-*Cas9*

**Table S10**: RNAseq differential expression analysis comparing Vasa-*Cas9 /*gFLE(G) transheterozygotes vs. gFLE(G)

**Table S11:** Gene ontology (GO) enrichment analysis using the significantly down-regulated genes in three Cas9/gRNA vs control comparisons.

**Table S12**: Gene ontology (GO) enrichment analysis using the significantly up-regulated genes in three Cas9/gRNA vs control comparisons.

**Table S13:** Mendelian ratios of one day old larvae from +/gFLE X +/Cas9 -females survive embryogenesis.

**Table S14:** Survival of genotypes through larval stage -females die before pupation.

**Table S15:** gFLE/Cas9 adult male survival curves

**Table S16:** gFLE/Cas9 males bulk percent fertility.

**Table S17:** Raw sex-genotype pupae counts of F1 offspring from homozygous gFLE_G_/gFLE_G_ males X Cas9/Cas9 females. (Presented in **Figure 1E**)

**Table S18** : Raw sex-genotype pupae count for **F2** offspring from +/gFLE X +/Cas9 crosses. (Presented in; **Fig 1D** Bottom row left (gFLE_G_), and **Fig S8** (gFLE_I_, gFLE_J_)

**Table S19** : Raw sex-genotype pupae count for **F3** offspring from +/gFLE X +/Cas9 crosses. (Presented in; **Fig 1D** Bottom row right (gFLE_G_), and **Fig S8** (gFLE_I_, gFLE_J_)

**Table S20:** Parameters used in *Anopheles gambiae* population suppression model.

**Table S21:** All Primers and gBlocks used in this study.

## SUPPLEMENTARY FIGURES

**Fig S1: System summary**

**Fig S2: gRNA Mutation summary Fig S3: Y chrom PCRs**

**Fig S4: Nanopore/RNA Seq**

**Fig S5: RNAseq Sample clustering Fig S6: Female embryo survival**

**Fig S7: Female larval lethality assay Fig S8: Male survival assay**

**Fig S9: F2s and F3s of PPs from gFLEI and gFLEJ Fig S10: Frequency of Δfle mutations in F2 offspring**

## SUPPLEMENTARY FIGURES

**Figure S1.**
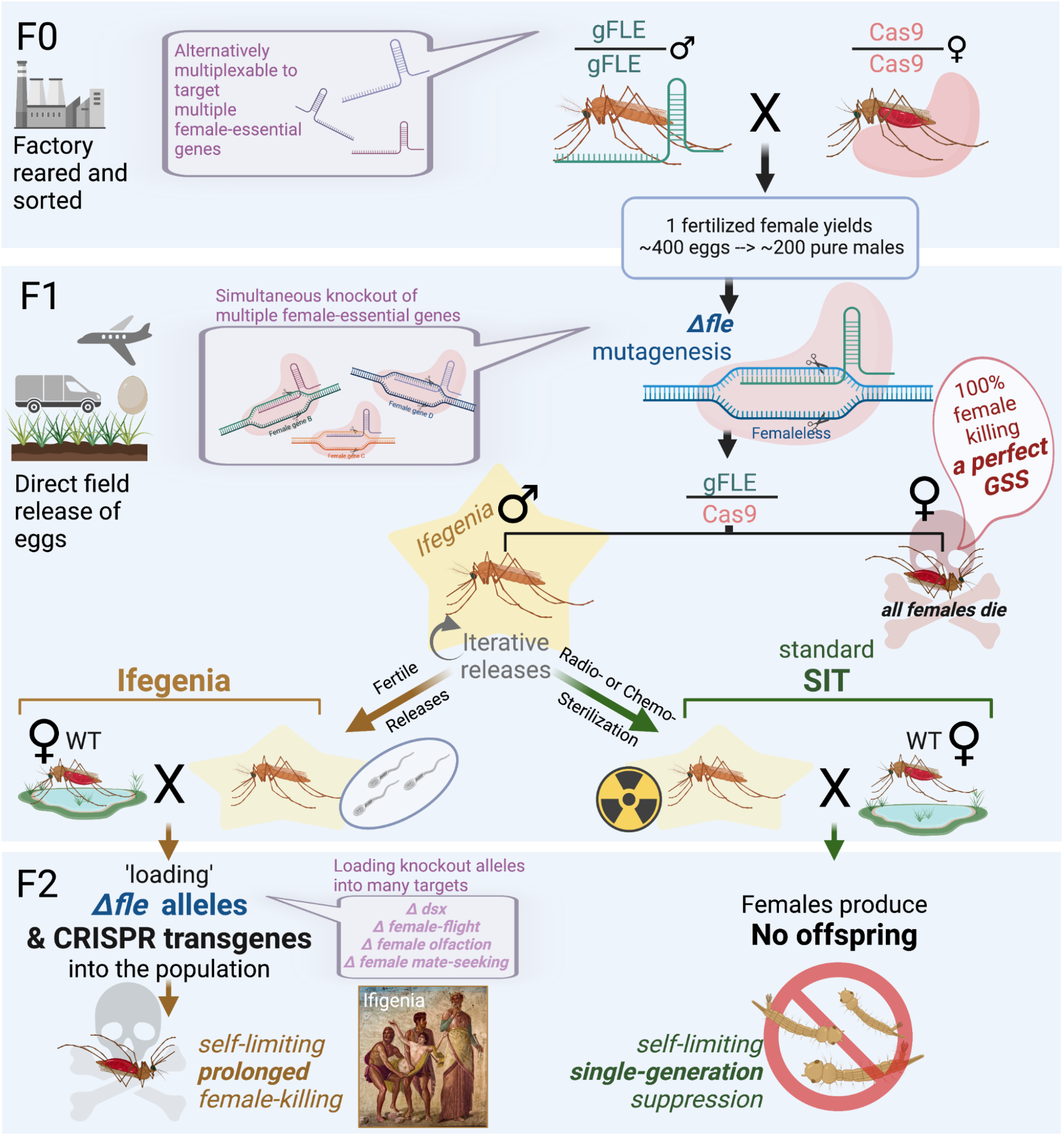
Using Ifegenia males for vector control. **F0)** Mass, factory-based, rearing and sorting of males from a stock gFLE/gFLE line, and females from a Cas9/Cas9 line yields a F0 cross from which each fertilized female can produce ∼200 Ifegenia sons during the course of her lifetime. (multiplexed iteration targeting multiple female-essential targets shown in adjacent bubbles, purple). **F1)** Produced F1 eggs undergo significant *Δfle* mutagenesis are expected to be 100% Ifegenia males, producing a perfect Genetic Sexing Strain (GSS). Therefore these eggs can be directly released into the environment as part of an Ifegenia multi-generational suppression system (left), or sterilized for release as part of a more traditional SIT-based system (right). Once grown, the resulting F1 males will mate with wild type females to produce their respective population-suppression effects. **F2)** In a multi-generational suppression system, Ifegenia males remain fertile but ‘load’ *Δfle* alleles into the population to cause prolonged suppression by daughter killing - providing a non-driving technology. In a traditional SIT-based system involving male sterilization, no F2 offspring would result from released F1 Ifegenia males (right). Due to its profound daughter-killing phenotypes, we’ve termed it Ifegenia (**<u>I</u>**nherited **<u>F</u>**emale **<u>E</u>**limination by **<u>G</u>**enetically **<u>E</u>**ncoded **<u>N</u>**ucleolyticly **<u>I</u>**nterrupted **<u>A</u>**lleles), in honor of Iphigenia of Greek mythology who was sacrificed by her father, King Agamemnon, to win a great battle.

**Figure S2:**
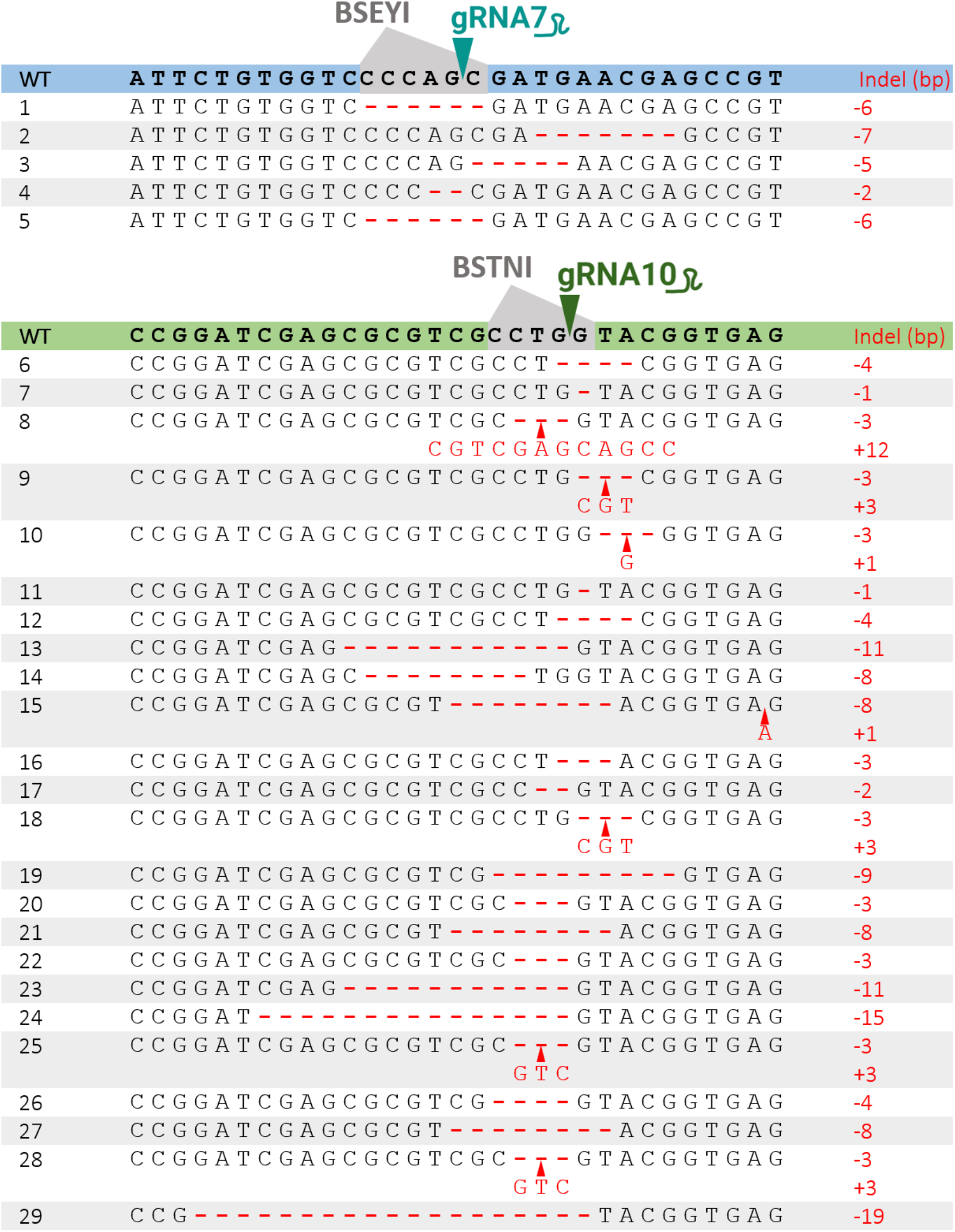
gFLE/Cas9 individuals have *Δfle* mutations under gRNA target sites. **gRNA7 (top):** Reads 1-5 are from a +_J_/+ larvae, gFLE_G_/Cas9 adult male, gFLE_J_/Cas9 larvae, a F2 gFLE_I_/Cas9 phenotypic female that survived to adulthood, and a F3 gFLE_J_/Cas9 intersex individual that died during eclosure respectively. **gRNA10 (bottom):** Reads 6-14 are from F2 larvae of mixed genotypes encompassing most genotypes and gFLE families. Read 15 was isolated from the only F1 gFLE_G_/Cas9 phenotypic female identified (**Table S6**), however which later PCR amplified for the presence of a Y-chromosome suggesting feminization. Reads 16 and 17 were isolated from a F2 gFLE_I_/Cas9 phenotypic female that survived to adulthood (**Table S4**), which also contained the gRNA7-derived mutation listed in Read 4. Reads 18 and 19 were identified from a F2 gFLE_J_/Cas9 phenotypic female that died as a pupae. Reads 20 and 21 are from a F2 gFLE_J_/Cas9 female that died as an adult. Reads 22 and 23 are from a F3 gFLE_J_/Cas9 intersex individual that died during eclosure, which also contained gRNA7-derived Read 5. Reads 24-29 are from gFLE_G_ /Cas9 females derived from paternally-derived Cas9 which are not otherwise discussed in this work.

**Figure S3.**
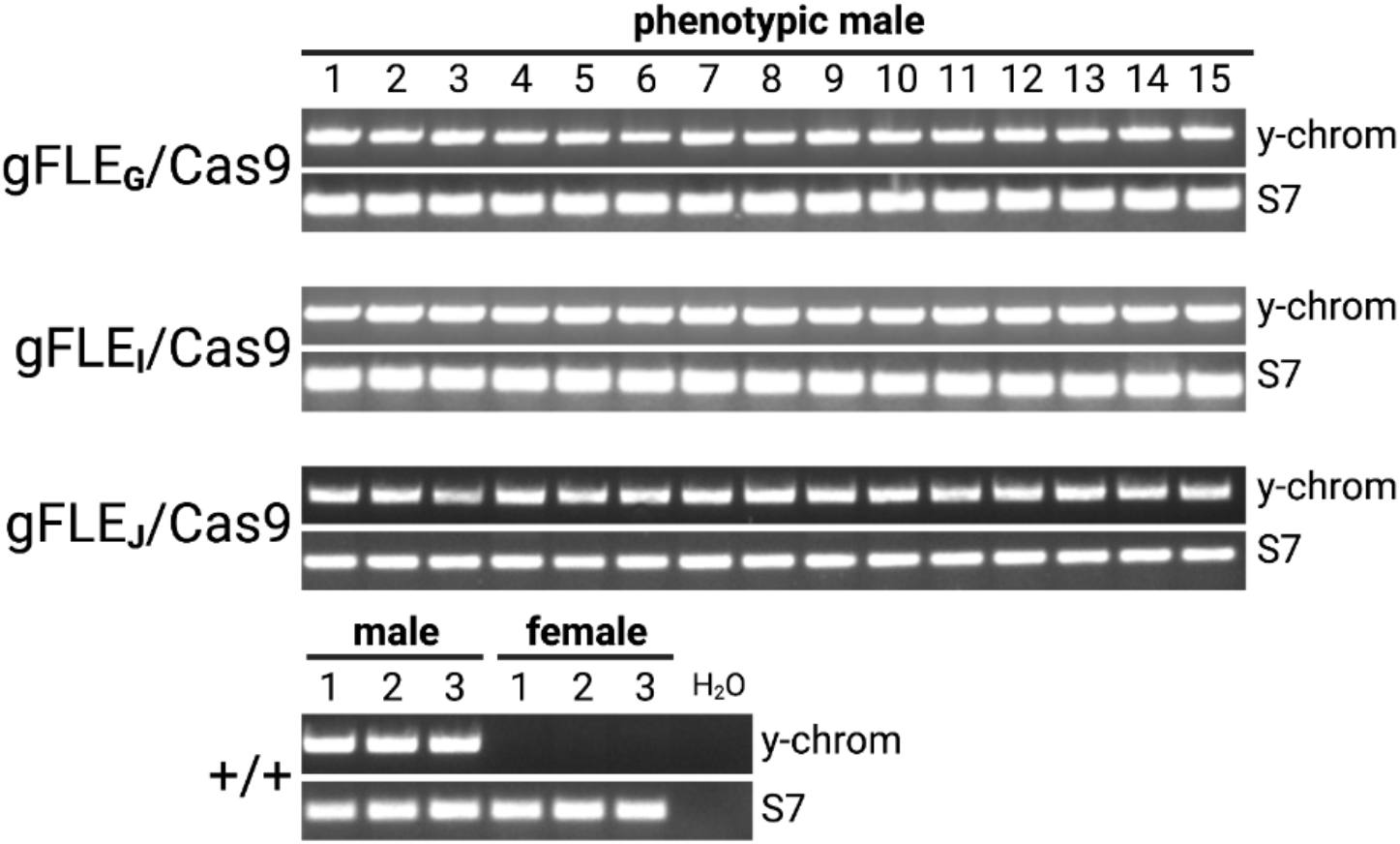
A random selection of gFLE/Cas9 transheterozygous adults which appeared phenotypically male were PCR amplified for the presence of the Y-chromosome.

**Figure S4.**
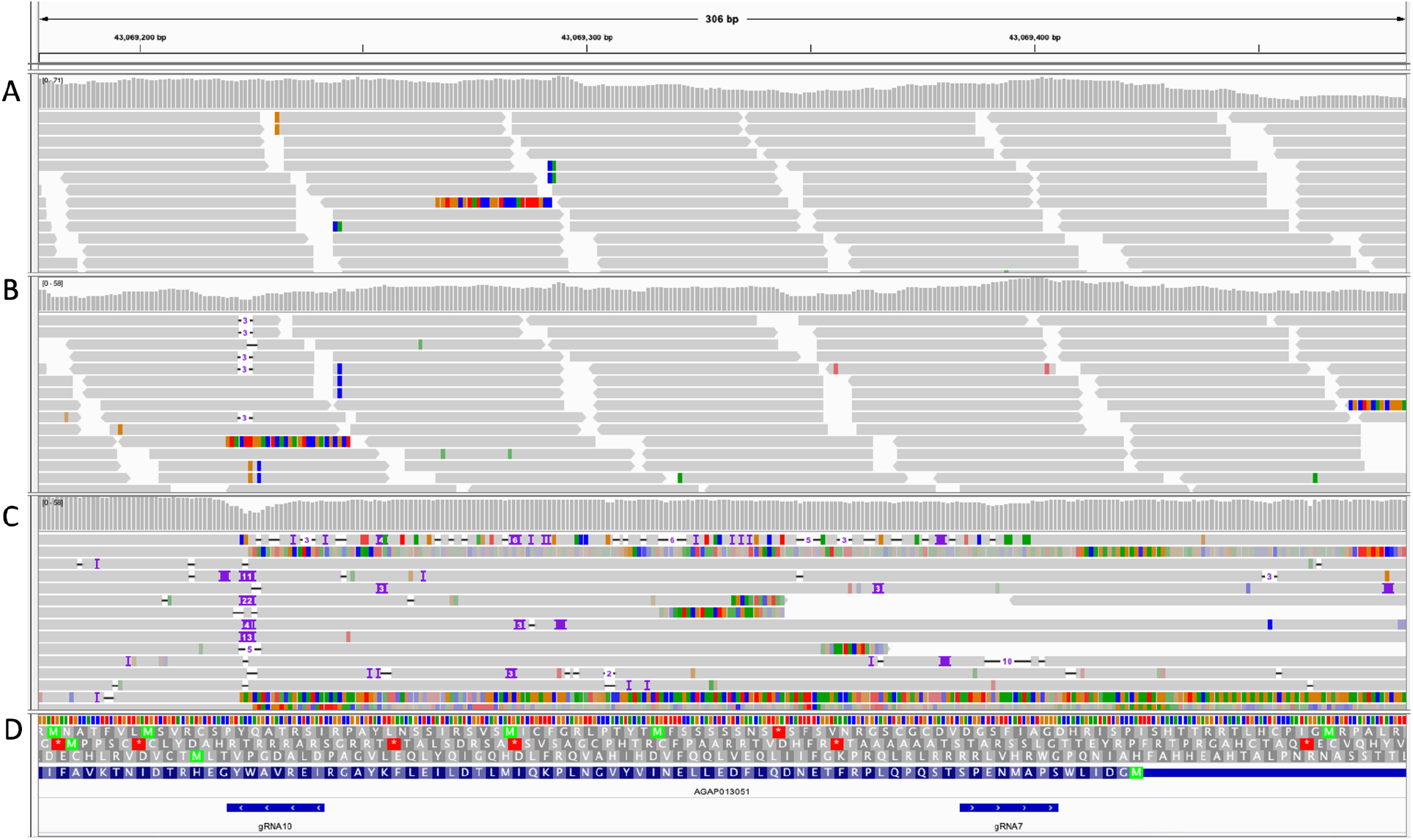
Genome browser snapshot of the *fle* gene (AGAP013051) zooming in on the gRNA target regions (2R:43,067,178-43,069,484). **A)** RNAseq reads of the WT control embryos. **B)** RNAseq reads of the gRNA/Cas9 transheterozygous embryos. **C)** Nanopore DNA sequencing of gRNA/Cas9 transheterozygous adult males. **D)** gRNA 10 and gRNA7 target sites indicated by blue bars.

**Figure S5.**
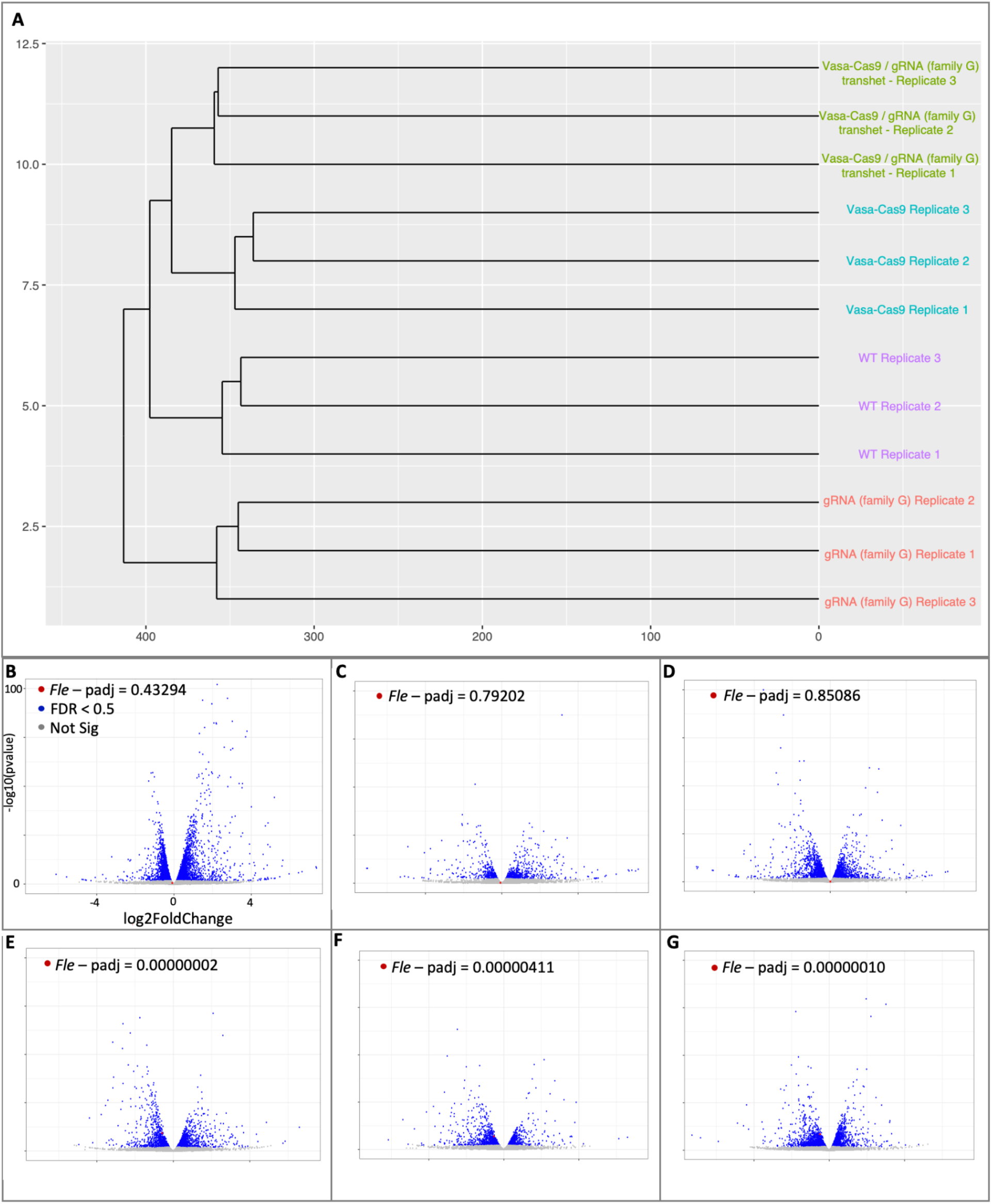
RNAseq samples clustering and volcano plots. **A)** Hierarchical clustering of the 12 RNAseq samples. Volcano plots comparing: **B)** gRNA(Family G) replicates to the WT control replicates; **C)** Vasa-Cas9 replicates to the WT control replicates; **D)** Vasa-Cas9 replicates to the gRNA(Family G) replicates; **E)** Vasa-Cas9**/**gRNA(Family G) transhet replicates to WT control replicates; **F)** Vasa-Cas9**/**gRNA(Family G) transhet replicates to Vasa-Cas9 replicates; **G)** Vasa-Cas9**/**gRNA(Family G) transhet replicates to gRNA(Family G) replicates. The X and Y axis are the same for plots B-G.

**Figure S6.**
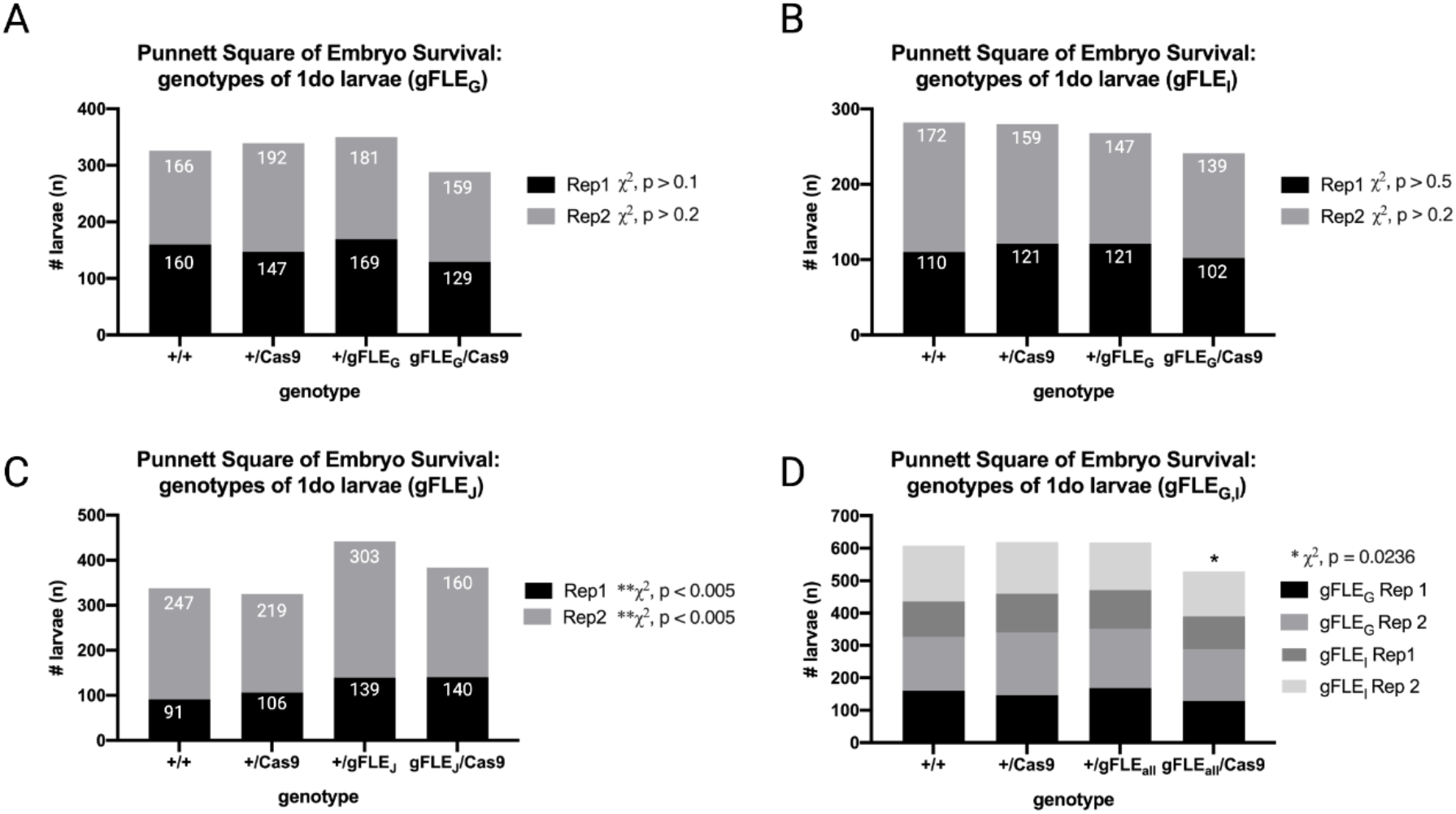
Mosaic mutant *Δfle* females survive embryogenesis. Reported as 1do larvae counts by genotype of the offspring from a cross of gFLE/+ males to Cas9/+ females. Significant embryonic female death would be expected to be observed as an approximate halving of the gFLE/Cas9 group. **A)** Raw counts of 1do larvae by genotype, offspring from a +/gFLE_G_ **♂** to +/Cas9♀cross. Two replicates shown stacked in black and grey. Both not significantly different from expected 1:1:1:1 Mendelian ratios, (χ^2^, p > 0.1 and p > 0.2 respectively). **B)** 1do larvae counts from a +/gFLE_I_**♂** to +/Cas9♀ cross. Two replicates shown stacked in black and grey. Both are not significantly different from expected 1:1:1:1 Mendelian ratios, (χ^2^, p > 0.5 and p > 0.2 respectively). **C)** 1do larvae counts from a +/gFLE_J_**♂** to +/Cas9♀ cross. Two replicates shown stacked in black and grey. Both significantly different from expected 1:1:1:1 Mendelian ratios (both χ^2^, p < 0.005) consistent with multiple insertions of transgene gFLE_J._ Family gFLE_J_ was therefore omitted from the analysis in **D). D)** Data pooled from **A)** and **B)**. Together these results demonstrate very slight levels of female embryo lethality in the gFLE/Cas9 group (χ^2^ p = 0.0236)

**Figure S7.**
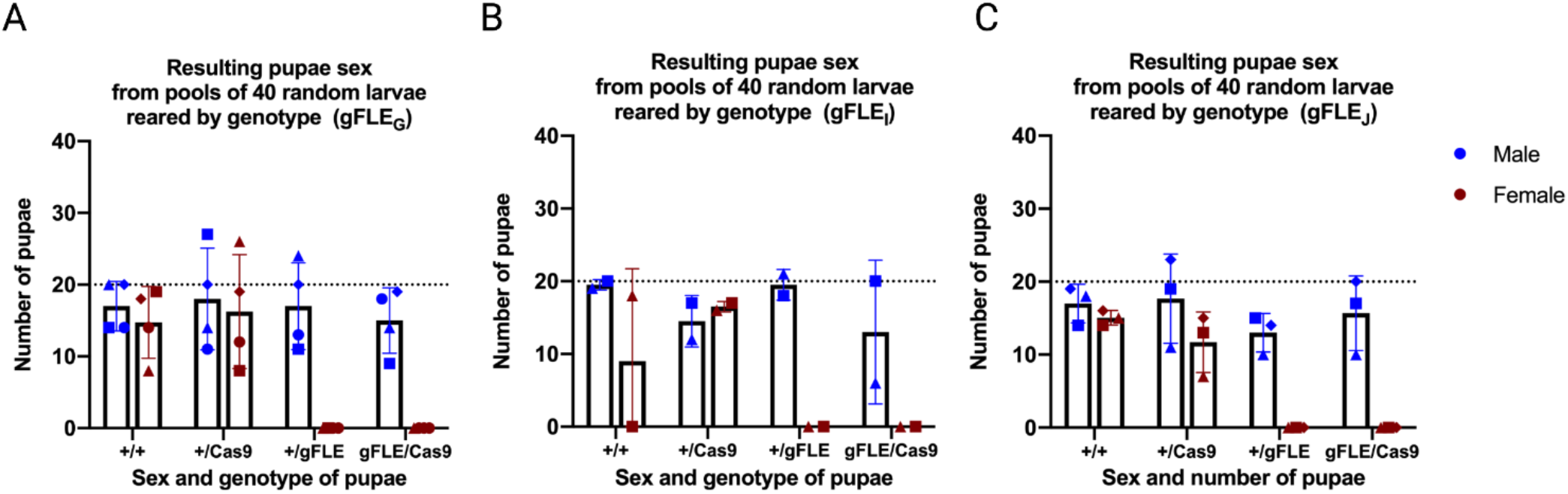
*Femaleless* mosaic Δ*fle* mutants (gFLE/+ and gFLE/Cas9) die during larvaehood. 40 random 1 day old larvae were isolated into trays by genotype and reared separately. The number, sex, and genotype of individuals is reported upon pupation. Replicates 1-4 are denoted by triangle, square, diamond, and circle respectively **A)** Δ*fle* females (gFLE_G_/+ and gFLE_G_/Cas9) were present at 1do but failed to pupate, all 4 replicates shown. **B)** Δ*fle* females (gFLE_I_/+ and gFLE_I_/Cas9) were present at 1do but failed to pupate. A third replicate was not performed on this line as the line was deemed sub-optimal for release and omitted from downstream analysis. **C)** Δ*fle* females (gFLE_J_/+ and gFLE_J_/Cas9) from family were present at 1do but failed to pupate, three replicates shown.

**Supplementary Figure S8.**
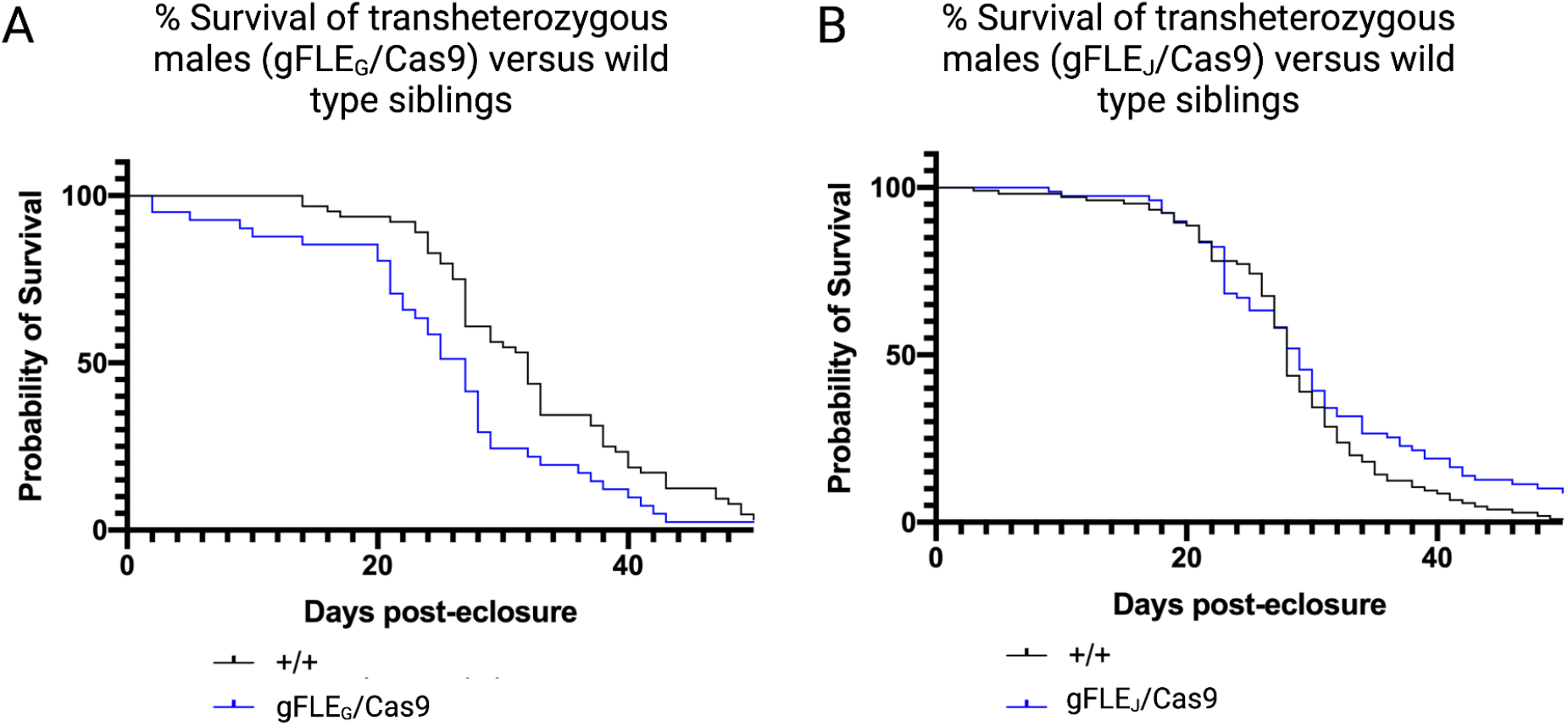
Male gFLE/Cas9 transheterozygous die only slightly faster than wild type siblings. **A)** gFLE_G_/Cas9 transheterozygous males (n = 40) die faster than wild type siblings (N = 74) (Log-rank p = 0.0017) **B)** gFLE_J_/Cas9 trans heterozygous males (n = 105) die slightly faster than wild type siblings (n = 79) (Log-rank, p= 0.0197).

**Supplementary Figure S9:**
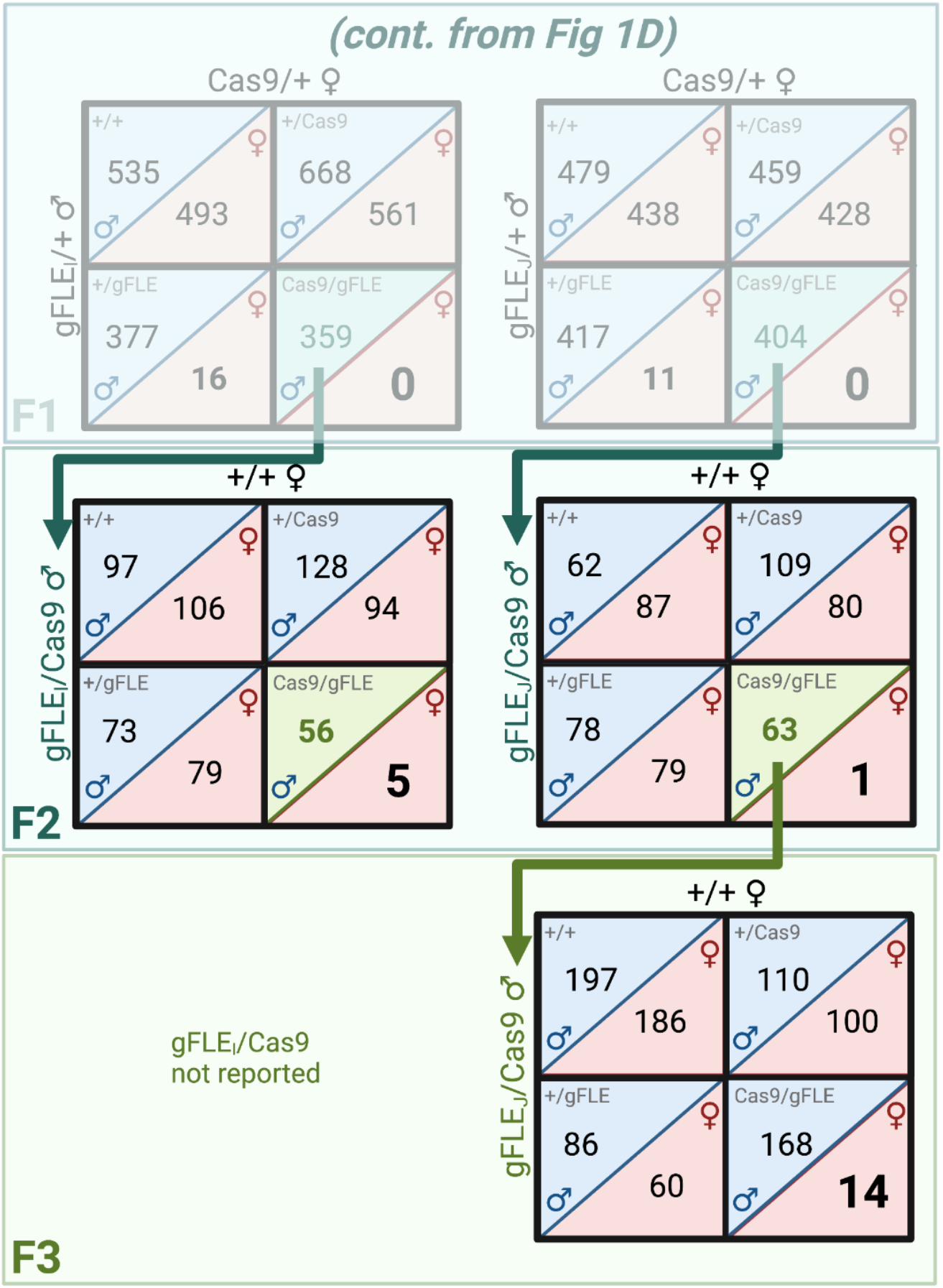
Female killing persists in F2 and F3 generations gFLE_I_/Cas9 and gFLE_J_/Cas9. In multigenerational experiments following from the crosses outlined in **Figure 1C**, F2 offspring from family gFLE_I_ were followed through the F2 generation, and offspring from family gFLE_J_ were followed through the F3 generation. Phenotypic gFLE/Cas9 females were identified in both families though at significantly reduced frequencies than should be expected by Mendelian segregation.

**Supplementary Figure 10:**
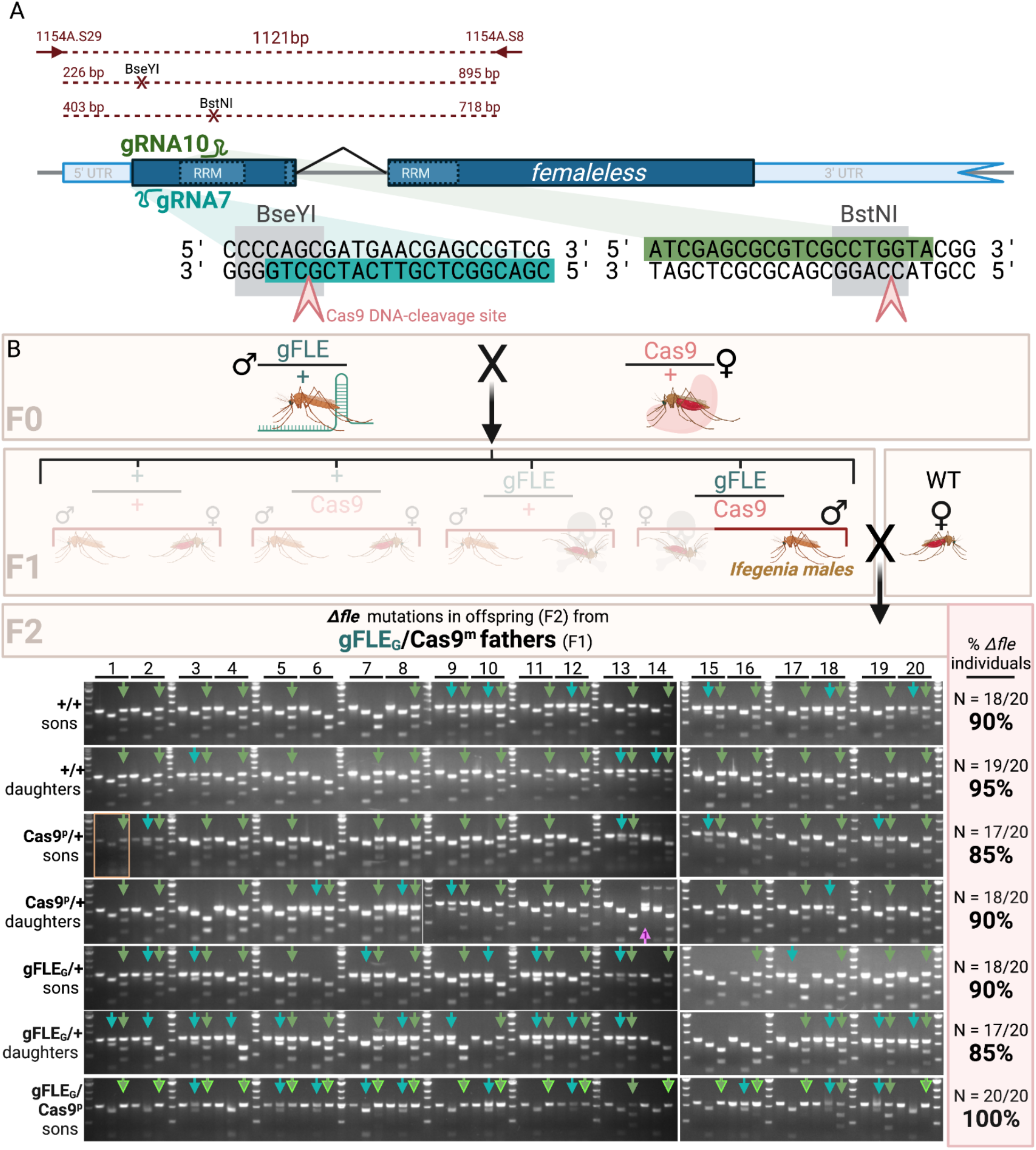
Approximate frequency of *Δfle* alleles in F2 offspring of gFLE_G_/Cas9^m^(F1) males. Hybrid F1 males inherited Cas9 maternally (Cas9^m^) **A)** To quantify approximate knockout frequencies under the gRNA7 and gRNA 10 target sites, *fle* was PCR-amplified with primers 1154A.S29 and 1154A.S8 (red) and subsequently digested with either restriction enzyme BseYI (gRNA7) or BstNI (gRNA10). These enzymes have semi-unique recognition sites (gray boxes) in wild type overlapping the predicted CRISPR mutagenesis sites (salmon chevron). CRISPR mutagenesis will preclude enzyme digestion of the PCR amplicons corresponding to that allele for most mutations. In diploids, PCR of wild type sequences should fully digest, PCR of heterozygotes should partially digest, and PCR of homozygotes should fail to digest. **B)** 20 gFLE_G_/Cas9 individuals for each genotype-sex are shown numbered across the top in brackets. Three consecutive wells are loaded for each sample, left to right: undigested *fle* PCR product, BseYI-digested PCR product, BstNI-digested PCR product. Undigested BseYI and BstNI bands indicate probable CRISPR mutations under gRNA 7 (teal arrows) and gRNA 10 (olive arrows), respectively. Biallelic Δ*fle* mutants are noted (highlighted arrows), most of which are in the gFLE_G_/Cas9 group due to active mosaic mutagenesis. Probable large insertions or deletions are denoted with purple arrows. Areas of the gel where exposure and brightness were adjusted separately are noted with orange boxes. The frequency of *Δfle* mutant individuals (those with at least one mutation) in that genotype-sex cohort are summarized at right in yellow.

## References

1. W. H. Organization, Others, World malaria report 2020: 20 years of global progress and challenges (2020) (available at https://apps.who.int/iris/bitstream/handle/10665/337660/9789240015791-eng.pdf).

2. H. Ranson, N. Lissenden, Insecticide Resistance in African Anopheles Mosquitoes: A Worsening Situation that Needs Urgent Action to Maintain Malaria Control. Trends Parasitol. 32, 187–196 (2016).

3. S. Sougoufara, E. C. Ottih, F. Tripet, The need for new vector control approaches targeting outdoor biting Anopheline malaria vector communities. Parasit. Vectors. 13, 295 (2020).

4. P. A. Papathanos, K. Bourtzis, F. Tripet, H. Bossin, J. F. Virginio, M. L. Capurro, M. C. Pedrosa, A. Guindo, L. Sylla, M. B. Coulibaly, F. A. Yao, P. S. Epopa, A. Diabate, A perspective on the need and current status of efficient sex separation methods for mosquito genetic control. Parasit. Vectors. 11, 654 (2018).

5. M. Zacarés, G. Salvador-Herranz, D. Almenar, C. Tur, R. Argilés, K. Bourtzis, H. Bossin, I. Pla, Exploring the potential of computer vision analysis of pupae size dimorphism for adaptive sex sorting systems of various vector mosquito species. Parasit. Vectors. 11, 656 (2018).

6. H. Yamada, M. J. B. Vreysen, K. Bourtzis, W. Tschirk, D. D. Chadee, J. R. L. Gilles, The Anopheles arabiensis genetic sexing strain ANO IPCL1 and its application potential for the sterile insect technique in integrated vector management programmes. Acta Trop. 142, 138– 144 (2015).

7. M. L. Taracena, C. M. Hunt, M. Q. Benedict, P. M. Pennington, E. M. Dotson, Downregulation of female doublesex expression by oral-mediated RNA interference reduces number and fitness of Anopheles gambiae adult females. Parasit. Vectors. 12, 170 (2019).

8. F. Bernardini, R. Galizi, M. Menichelli, P.-A. Papathanos, V. Dritsou, E. Marois, A. Crisanti, N. Windbichler, Site-specific genetic engineering of the Anopheles gambiae Y chromosome. Proc. Natl. Acad. Sci. U. S. A. 111, 7600–7605 (2014).

9. F. Catteruccia, J. P. Benton, A. Crisanti, An Anopheles transgenic sexing strain for vector control. Nat. Biotechnol. 23, 1414–1417 (2005).

10. A. L. Smidler, S. N. Scott, E. Mameli, W. R. Shaw, F. Catteruccia, A transgenic tool to assess Anopheles mating competitiveness in the field. Parasit. Vectors. 11, 651 (2018).

11. E. Marois, C. Scali, J. Soichot, C. Kappler, E. A. Levashina, F. Catteruccia, High-throughput sorting of mosquito larvae for laboratory studies and for future vector control interventions. Malaria Journal. 11 (2012),, doi:10.1186/1475-2875-11-302.

12. G. Fu, R. S. Lees, D. Nimmo, D. Aw, L. Jin, P. Gray, T. U. Berendonk, H. White-Cooper, S. Scaife, H. Kim Phuc, O. Marinotti, N. Jasinskiene, A. A. James, L. Alphey, Female-specific flightless phenotype for mosquito control. Proc. Natl. Acad. Sci. U. S. A. 107, 4550–4554 (2010).

13. M. Li, T. Yang, M. Bui, S. Gamez, T. Wise, N. P. Kandul, J. Liu, L. Alcantara, H. Lee, J. R. Edula, R. Raban, Y. Zhan, Y. Wang, N. DeBeaubien, J. Chen, H. M. Sánchez C, J. B. Bennett, I. Antoshechkin, C. Montell, J. M. Marshall, O. S. Akbari, Suppressing mosquito populations with precision guided sterile males. Nat. Commun. 12, 5374 (2021).

14. R. Galizi, L. A. Doyle, M. Menichelli, F. Bernardini, A. Deredec, A. Burt, B. L. Stoddard, N. Windbichler, A. Crisanti, A synthetic sex ratio distortion system for the control of the human malaria mosquito. Nat. Commun. 5, 3977 (2014).

15. F. A. Yao, A.-A. Millogo, P. S. Epopa, A. North, F. Noulin, K. Dao, M. Drabo, C. Guissou, S. Kekele, M. Namountougou, R. K. Ouedraogo, L. Pare, N. Barry, R. Sanou, H. Wandaogo, R. K. Dabire, A. McKemey, F. Tripet, A. Diabaté, Mark-release-recapture experiment in Burkina Faso demonstrates reduced fitness and dispersal of genetically-modified sterile malaria mosquitoes. Nat. Commun. 13, 796 (2022).

16. E. Krzywinska, J. Krzywinski, Effects of stable ectopic expression of the primary sex determination gene Yob in the mosquito Anopheles gambiae. Parasit. Vectors. 11, 648 (2018).

17. K. Kyrou, A. M. Hammond, R. Galizi, N. Kranjc, A. Burt, A. K. Beaghton, T. Nolan, A. Crisanti, A CRISPR–Cas9 gene drive targeting doublesex causes complete population suppression in caged Anopheles gambiae mosquitoes. Nat. Biotechnol. 36, 1062–1066 (2018).

18. K. C. Long, L. Alphey, G. J. Annas, C. S. Bloss, K. J. Campbell, J. Champer, C.-H. Chen Choudhary, G. M. Church, J. P. Collins, K. L. Cooper, J. A. Delborne, O. R. Edwards, C. I. Emerson, K. Esvelt, S. W. Evans, R. M. Friedman, V. M. Gantz, F. Gould, S. Hartley, E. Heitman, J. Hemingway, H. Kanuka, J. Kuzma, J. V. Lavery, Y. Lee, M. Lorenzen, J. E. Lunshof, J. M. Marshall, P. W. Messer, C. Montell, K. A. Oye, M. J. Palmer, P. A. Papathanos, P. N. Paradkar, A. J. Piaggio, J. L. Rasgon, G. Rašić, L. Rudenko, J. R. Saah, M. J. Scott, J. T. Sutton, A. E. Vorsino, O. S. Akbari, Core commitments for field trials of gene drive organisms. Science. 370, 1417–1419 (2020).

19. A. K. Beaghton, A. Hammond, T. Nolan, A. Crisanti, A. Burt, Gene drive for population genetic control: non-functional resistance and parental effects. Proc. Biol. Sci. 286, 20191586 (2019).

20. E. Krzywinska, L. Ferretti, J. Li, J.-C. Li, C.-H. Chen, J. Krzywinski, femaleless Controls Sex Determination and Dosage Compensation Pathways in Females of Anopheles Mosquitoes. Current Biology. 31 (2021), pp. 1084–1091.e4.

21. K. Werling, W. R. Shaw, M. A. Itoe, K. A. Westervelt, P. Marcenac, D. G. Paton, D. Peng, N. Singh, A. L. Smidler, A. South, A. A. Deik, L. Mancio-Silva, A. R. Demas, S. March, E. Calvo, S. N. Bhatia, C. B. Clish F. Catteruccia, Steroid Hormone Function Controls Non-competitive Plasmodium Development in Anopheles. Cell. 177, 315–325.e14 (2019).

22. G. M. Chambers, M. J. Klowden, Age of Anopheles gambiae Giles male mosquitoes at time of mating influences female oviposition. J. Vector Ecol. 26, 196–201 (2001).

23. Sawadogo, Diabaté, Toé, Effects of Age and Size on Anopheles gambiae ss Male Mosquito Mating Success. J. Med. Surg. Pathol. (available at https://academic.oup.com/jme/article-abstract/50/2/285/853180).

24. A. L. Smidler, O. Terenzi, J. Soichot, E. A. Levashina, E. Marois, Targeted mutagenesis in the malaria mosquito using TALE nucleases. PLoS One. 8, e74511 (2013).

25. N. P. Kandul, J. Liu, H. M. Sanchez C, S. L. Wu, J. M. Marshall, O. S. Akbari, Transforming insect population control with precision guided sterile males with demonstration in flies. Nat. Commun. 10, 84 (2019).

26. H. M. S. C., H. M. Sánchez C., S. L. Wu, J. B. Bennett, J. M. Marshall, MGD riv E: A modular simulation framework for the spread of gene drives through spatially explicit mosquito populations. Methods in Ecology and Evolution. 11 (2020), pp. 229–239.

27. D. O. Carvalho, A. R. McKemey, L. Garziera, R. Lacroix, C. A. Donnelly, L. Alphey, A. Malavasi, M. L. Capurro, Suppression of a Field Population of Aedes aegypti in Brazil by Sustained Release of Transgenic Male Mosquitoes. PLoS Negl. Trop. Dis. 9, e0003864 (2015).

28. J.-M. O. Depinay, C. M. Mbogo, G. Killeen, B. Knols, J. Beier, J. Carlson, J. Dushoff, P. Billingsley, H. Mwambi, J. Githure, A. M. Toure, F. E. McKenzie, A simulation model of African Anopheles ecology and population dynamics for the analysis of malaria transmission. Malar. J. 3, 29 (2004).

29. E. Krzywinska, N. J. Dennison, G. J. Lycett, J. Krzywinski, A maleness gene in the malaria mosquito Anopheles gambiae. Science. 353, 67–69 (2016).

30. R. Galizi, A. Hammond, K. Kyrou, C. Taxiarchi, F. Bernardini, S. M. O’Loughlin, P.-A. Papathanos, T. Nolan, N. Windbichler, A. Crisanti, A CRISPR-Cas9 sex-ratio distortion system for genetic control. Sci. Rep. 6, 31139 (2016).

31. G. Volohonsky, O. Terenzi, J. Soichot, D. A. Naujoks, T. Nolan, N. Windbichler, D. Kapps, L. Smidler, A. Vittu, G. Costa, S. Steinert, E. A. Levashina, S. A. Blandin, E. Marois, Tools for Anopheles gambiae Transgenesis. G3. 5, 1151–1163 (2015).

32. A. Hammond, R. Galizi, K. Kyrou, A. Simoni, C. Siniscalchi, D. Katsanos, M. Gribble, D. Baker, E. Marois, S. Russell, A. Burt, N. Windbichler, A. Crisanti, T. Nolan, A CRISPR-Cas9 gene drive system targeting female reproduction in the malaria mosquito vector Anopheles gambiae. Nat. Biotechnol. 34, 78–83 (2016).

33. K. M. Esvelt, A. L. Smidler, F. Catteruccia, G. M. Church, Concerning RNA-guided gene drives for the alteration of wild populations. Elife. 3 (2014), doi:10.7554/eLife.03401.

34. J. Champer, A. Buchman, O. S. Akbari, Cheating evolution: engineering gene drives to manipulate the fate of wild populations. Nat. Rev. Genet. 17, 146–159 (2016).

35. N. P. Kandul, J. Liu, J. B. Bennett, J. M. Marshall, O. S. Akbari, A confinable home-and-rescue gene drive for population modification. Elife. 10 (2021), doi:10.7554/eLife.65939.

36. A. Simoni, A. M. Hammond, A. K. Beaghton, R. Galizi, C. Taxiarchi, K. Kyrou, D. Meacci, M. Gribble, G. Morselli, A. Burt, T. Nolan, A. Crisanti, A male-biased sex-distorter gene drive for the human malaria vector Anopheles gambiae. Nat. Biotechnol. 38, 1054– 1060 (2020).

37. F. Sánchez-Bayo, K. A. G. Wyckhuys, Worldwide decline of the entomofauna: A review of its drivers. Biological Conservation. 232 (2019), pp. 8–27.

38. H. van den Berg, H. S. da Silva Bezerra, S. Al-Eryani, E. Chanda, B. N. Nagpal, T. B. Knox, R. Velayudhan, R. S. Yadav, Recent trends in global insecticide use for disease vector control and potential implications for resistance management. Sci. Rep. 11, 23867 (2021).

39. M. Q. Benedict, The MR4 Methods in Anopheles research laboratory manual. Atlanta: CDC; 2007 (2018).

40. Ho, Gesing, Consortium, VectorBase: an updated bioinformatics resource for invertebrate vectors and other organisms related with human diseases. Nucleic Acids Mol. Biol. (available at https://academic.oup.com/nar/article-abstract/43/D1/D707/2437683).

41. A. Smidler, thesis (2019).

42. J. Krzywinski, D. R. Nusskern, M. K. Kern, N. J. Besansky, Isolation and characterization of Y chromosome sequences from the African malaria mosquito Anopheles gambiae. Genetics. 166, 1291–1302 (2004).

43. R. J. Kent, A. J. West, D. E. Norris, Molecular differentiation of colonized human malaria vectors by 28S ribosomal DNA polymorphisms. Am. J. Trop. Med. Hyg. 71, 514–517 (2004).

44. A. Deredec, H. C. J. Godfray, A. Burt, Requirements for effective malaria control with homing endonuclease genes. Proc. Natl. Acad. Sci. U. S. A. 108, E874–80 (2011).

45. H. Li, Minimap2: pairwise alignment for nucleotide sequences. Bioinformatics. 34, 3094– 3100 (2018).

## Supplemental References

1. H. M. S. C., H. M. Sánchez C., S. L. Wu, J. B. Bennett, J. M. Marshall, MGD riv E: A modular simulation framework for the spread of gene drives through spatially explicit mosquito populations. Methods in Ecology and Evolution. 11 (2020), pp. 229–239.

